# Comparative metagenomics using pan-metagenomic graphs

**DOI:** 10.1101/2025.11.24.690211

**Authors:** Izaak Coleman, Natalya Mametyarova, Andrey Zaznaev, Peiwen Cai, Lisa Yu, Yoli Meydan, Aviya Litman, Ayushi Sharma, Lily He, Amanda Simkhovich, Dwayne Seeram, Heekuk Park, Yael R. Nobel, Aya Brown Kav, Itsik Pe’er, Anne-Catrin Uhlemann, Tal Korem

## Abstract

Identifying microbial genomic factors underlying human phenotypes is a key goal of microbiome research. Sequence graphs are a highly effective tool for genome comparisons because they enable high-resolution de novo analyses that capture and contextualize complex genomic variation. However, applying sequence graphs to complex microbial communities remains challenging due to the scale and complexity of metagenomic data. Existing multi-sample sequence graphs used in these settings are highly complex, computationally expensive, less accurate than single-sample alternatives, and often involve arbitrary coarse-graining. Here, we present copangraph, a multi-sample sequence-graph-based analysis framework for comprehensive comparisons of genomic variation across metagenomes. Copangraph uses a novel homology-based graph, which provides both non-arbitrary, evolutionary-motivated grouping of sequences into the same node as well as flexibility in the scale of variation represented by the graph. Its construction relies on hybrid coassembly, a new coassembly approach in which single-sample graphs are first constructed separately and are then merged to create a multi-sample graph. We also present an algorithm that uses paired-end reads to improve detection of contiguous genomic regions, increasing accuracy. Our results demonstrate that copangraph captures sequence and variant information more accurately than alternative methods, provides graphs that are more suitable for comparative analysis than de Bruijn graphs, and is computationally tractable. We show that copangraph reflects meaningful metagenomic variation across diverse scenarios. Importantly, it enables significantly better performance than other metagenomic representations when predicting the gut colonization trajectories of Vancomycin-resistant Enterococcus. Our results underscore the value of our multi-sample, graph-based framework for comparative metagenomic analyses.

## Introduction

Detecting genomic differences associated with phenotypes of interest is key in investigating their underlying mechanisms. As such, impactful discoveries have relied on identifying genomic differences associated with divergent phenotypes among similar strains, such as a Shiga-toxin-containing prophage that increased the virulence of an enteroaggregative *Escherichia coli* strain^1^, or genomic similarities associated with similar phenotypes in different taxa, such as a plasmid-borne extended-spectrum β-lactamase^2^. This type of analysis, termed “comparative genomics”^3–8^, has informed numerous discoveries regarding the pangenomes of specific species and genetic elements. “Comparative metagenomics” broadens its scope to simultaneously analyze the genomes of multiple different microbes across complex microbial communities (the “pan-metagenome”)^4,9^. Doing so in a sensitive, precise, scalable, and reference-free manner could advance our genetic and mechanistic understanding of complex host-microbiome associations. Furthermore, it would alleviate challenges with studying isolates, which requires a priori knowledge of a clade of interest, the ability to culture them, and ensuring that they do not acquire additional mutations post-isolation^10^.

Comparative metagenomics, however, is challenging: microbiomes are complex and dynamic, with multiple coexisting strains that are similar but not identical^11–16^. Many approaches to studying genomic variation in the microbiome handle this complexity by working with a single genomic sequence (or a collection of contigs) per taxon, which could be obtained from public sources^17–20^ or de-novo assembly^21–24^. While such approaches revealed intriguing insights into genomic variability in the microbiome, they are limited by the underlying genomic representation: each genome represents a single potential version of the taxon, and a set of genomes does not, in itself, facilitate the analysis of genomic variability without additional processing^25^.

Sequence graphs have been increasingly used to represent genomic variability across specific phylogenetic clades^26–29^, including the human genome^25,30,31^. In these graphs, nodes typically represent genomic sequences, and edges indicate that these sequences are contiguous in some genome. Branches in sequence graphs capture variability in the genomic structure^9,32^, allowing the graph as a whole to represent complex variation events between large sets of genomes and capture the variable elements within their genomic context^33–35^. Sequence graphs have immense potential in the metagenomic setting^9^, yet challenges inherent to the data have limited their adoption. Unlike monocultures, each microbiome sample contains multiple bacterial species, with each species exhibiting substantial intra-species variability, sometimes reaching >50% of the variable (accessory) genes within a species^4–7,36^. Horizontal exchange of genetic material and the activity of mobile genetic elements further complicate the metagenomic structure^37–39^. The sequencing data used to profile metagenomes is complex and noisy, provides insufficient coverage for many strains, and is typically composed of sequencing reads that are too short to fully traverse complex and repetitive genomic regions. Finally, to enable comparative metagenomics, analyses need to be performed at scale and across multiple samples, which requires computational tractability. As a result of this complexity, coassembling a single pan-metagenomic sequence graph that represents a collection of samples has proven both intractably resource-intensive^40–42^ and potentially of lower quality compared to single-sample assemblies, due to the increased heterogeneity resulting from merging multiple samples^43,44^.

Two approaches have emerged to handle these challenges while still leveraging sequence graphs. The first approach is to construct single-sample metagenomic graphs and then compare their various properties across samples^45–47^. For example, a recent analysis constructed hundreds of single-metagenome MetaCarvel scaffold graphs of samples from different human body sites, and then compared variant rates and properties across them^47^. In this approach, the topology of each single-sample graph does not depend upon cross-sample genomic variability, making specific genomic comparisons across samples challenging. The second approach is to coassemble multiple samples, but coarse-grain the resulting sequence graph to reduce its complexity. For example, spacegraphcats^48^ allows for rapid querying of k-mer occurrence within “neighborhoods” in the graph, arbitrarily defined using r-dominating sets, and MetaFast^49^ reduces the graph to connected components at an arbitrary size range, primarily for calculating distances between microbiomes. While these approaches facilitate comparative metagenomics, the coarse-graining is arbitrary, and limits both systematic analysis and the interpretability of the findings.

Here we present copangraph, an efficient **co**mparative **pan**-metagenomic **graph** designed to comprehensively represent the genomes of multiple microbes across many samples. Copangraph is a homology-based graph that captures evolutionary relationships between sequences. It represents both within- and between-sample variability by recording the sample from which each sequence, node, and edge in the graph originated, and facilitates reference-free comparative analysis, allowing complex variation among taxa to be studied in a metagenomic context. Copangraph construction is based on hybrid coassembly, a novel conceptual approach for *de novo* metagenomic analysis, in which samples are first assembled separately and subsequently merged. We introduce three methodological advances: a contiguity detection algorithm for constructing more accurate single-sample sequence graphs; a graph merging procedure that produces a cross-sample graph; and homology group clustering, which collapses related genomic regions into the same node based on a tunable homology threshold. In multiple benchmarks using both long-read sequencing and simulations, we demonstrate that our approach provides a more accurate representation of both single-sample metagenomes and multi-sample pan-metagenomes compared to alternative methods, while requiring fewer resources than traditional coassembly. Furthermore, we demonstrate that copangraphs provide an informative representation of the pan-metagenome that is better suited for comparative analyses than de Bruijn graphs. Finally, we show that copangraph offers informative metagenomic representations across different clinical scenarios, facilitates improved prediction of trajectories of antibiotic-resistant colonization relative to other metagenomic representations, and detects specific variable regions associated with colonization persistence and clearance.

## Results

### Copangraph, a multi-sample comparative pan-metagenomic sequence graph

To extend sequence-graph analysis to the comparative metagenomic setting, we developed copangraph, a multi-sample sequence-graph that enables de-novo comparison of variable genomic content between metagenomes. We developed a method for copangraph construction from multiple short-read paired-end metagenomic samples that is scalable to hundreds of samples (**Methods**). A copangraph of human gut metagenomes sampled from different patients in a liver-transplant cohort^14,50^ (**Methods**; **Fig. 1a**) shows between-taxon genomic interactions (**Fig. 1b,c**) and within-taxon heterogeneity (**Fig. 1d,e**).

**Figure 1.**
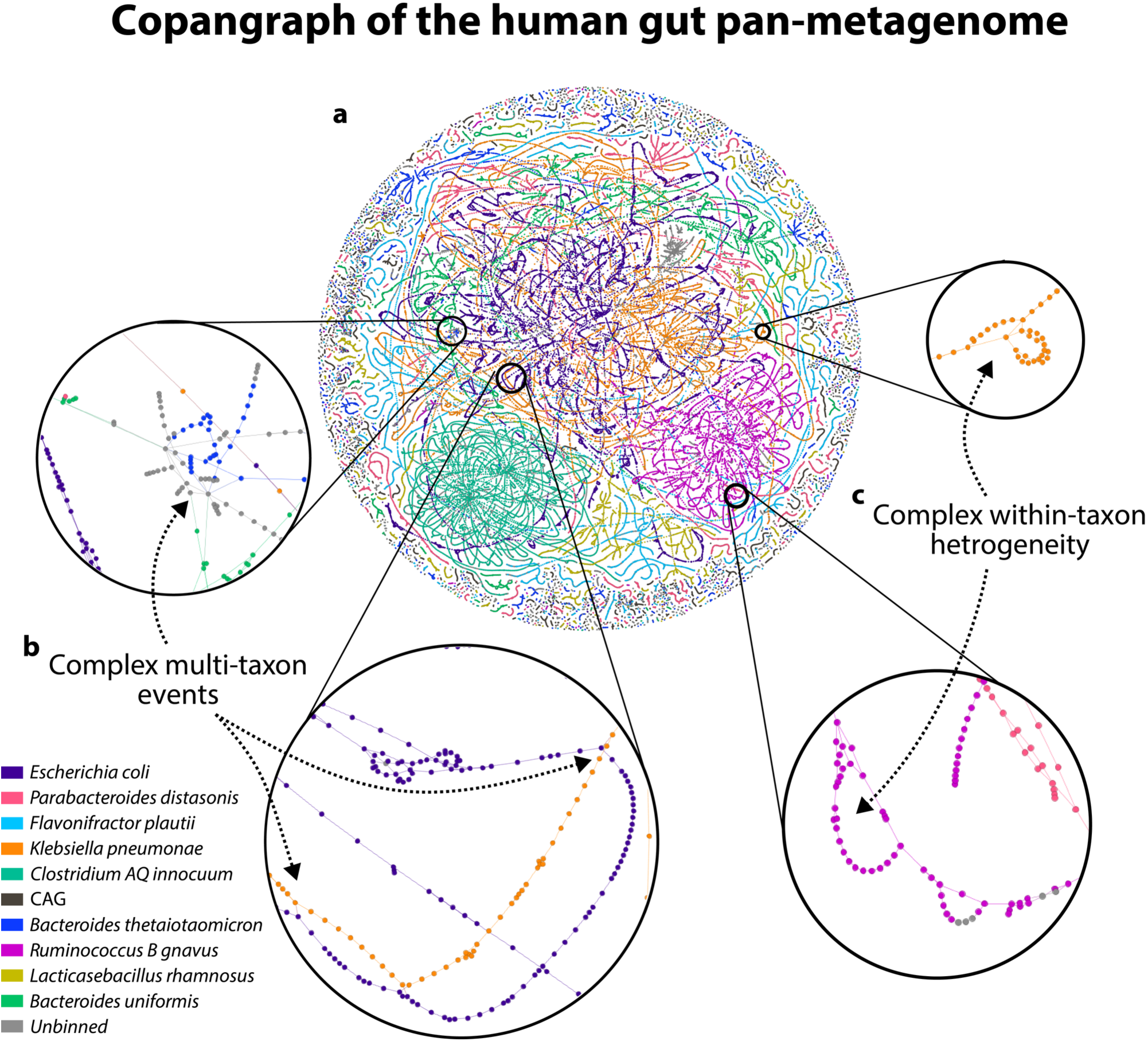
Copangraph captures complex sequence relationships among species of the human gut pan-metagenome. **a,** A copangraph constructed from gut metagenomic samples of liver transplant recipients (**Methods**). Shown are nodes colored according to species classification for the ten most frequent species (by number of nodes) as well as unclassified nodes (gray) that are at most five nodes from a classified node (**Methods**). **b,c,** Insets showing regions of the copangraph exhibiting close inter-species homologies (b) and putative intra-species heterogeneity (c). For display purposes, each colored point represents a 2500-bp region from a copangraph node.

### Overview of copangraph construction

Copangraph is built with an approach we term “hybrid coassembly”. Hybrid coassembly bridges traditional assembly efforts, in which each sample is assembled separately^23,44,51^, with traditional coassembly, in which reads from multiple metagenomic samples are pooled together and assembled as if they were a single sample^40^. In hybrid coassembly, each sample is first assembled separately to produce a sequence graph (**Fig. 2a**). Then, these single-sample graphs are merged together to create a multi-sample pan-metagenomic graph (**Fig. 2b**). This approach benefits from assembling each sample separately, which was shown to be more accurate than coassembly^43,44^, as well as from improved efficiency and parallelization.

**Figure 2.**
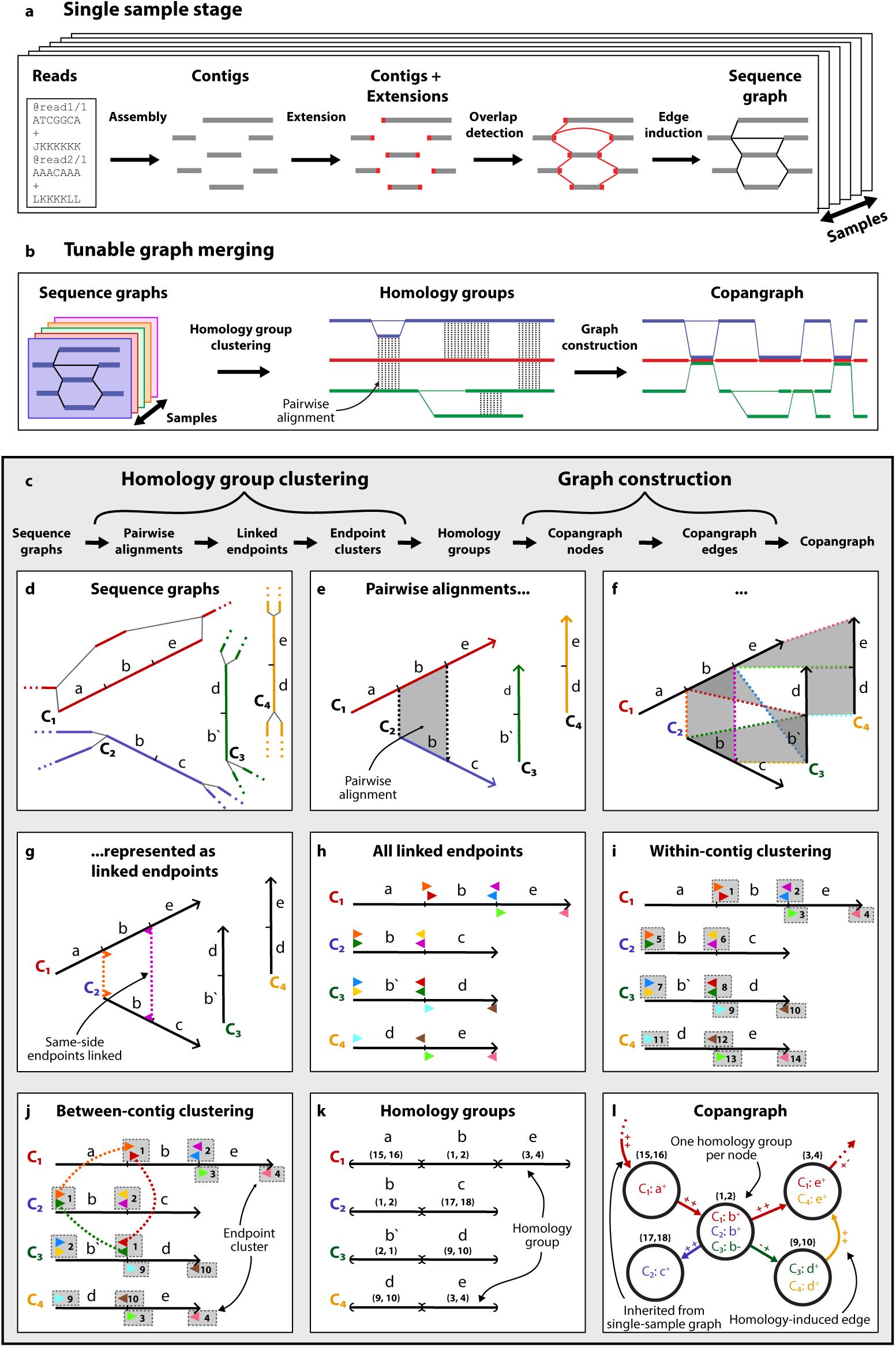
Constructing a multi-sample pan-metagenomic sequence graph. **a,b,** An overview of our hybrid coassembly approach: **a,** Samples are first processed separately, starting with assembly of each one into contigs. We then use paired-end reads to assemble the genomic region immediately contiguous to each contig (red boxes). Overlaps between these extensions and other contigs induce an edge (black line), creating a single-sample sequence graph. **b,** These sequence graphs are then merged to create a multi-sample pan-metagenomic graph. First, we identify groups of homologous sequences among the contigs present within each graph. Then, a node is created for each group, and edges are created whenever a contig or an edge in the sample-specific graphs connects between nodes. Note that in the final copangraph, the originating sample (color) of every node and edge is known. **c,** An in-depth explanation of tunable graph merging: **d,** Four sequence graphs colored by sample origin. The contigs C_1_-C_4_ are composed of intervals a-e, where similar letters indicate homology (and “b’” is homologous to “b” in the reverse complement). Note that in practice, homology is not yet known. **e,f,** We perform pairwise alignment between all contigs. **g,h,** To cluster these alignments, we represent each alignment as four linked endpoints. **i,** First, we cluster endpoints within each contig that point in the same direction and are in close proximity. Endpoints in the same cluster are shown in grey boxes with the same number (the cluster’s identity). **j,** We then cluster endpoints between contigs by merging within-contig clusters that share endpoints that are linked to one another due to their corresponding alignment (same color). **k,** Finally, we divide the contigs into disjoint intervals, each belonging to a homology group according to the cluster identity of the bounding endpoints. **l,** The intervals of each homology group are collapsed into a copangraph node. Bi-directed edges are induced between nodes if there was an edge between them in the single-sample sequence graph or if they contain sequences that are contiguous in one of the single-sample contigs. The bi-direction of an edge reflects the intervals’ orientations within the nodes it connects.

Single-sample assemblers optimize for assembly quality and contiguity, rather than for a graph representative of genomic structure^9,35^. Therefore, in the single-sample stage (**Fig. 2a**), we introduce a contiguity detection algorithm that recreates a sequence graph from assembled contigs: After assembling each sample, we assemble the genomic region immediately up- or downstream of each contig using an accurate and efficient extension algorithm that uses paired-end reads. We then detect overlaps between these extensions and other contigs, and use them to induce bi-directed edges (i.e., edges that reflect sequence orientation) in the resulting sequence graph (**Methods**).

Genomic variability occurs at multiple scales, from single-nucleotide polymorphisms (SNPs) to horizontal gene transfer or large-scale genomic rearrangements. This variation has a sizable effect on the fragmentation of contigs and the complexity of sequence graphs^52^. Many sequence-graph construction methods do not allow direct control over the level of variation represented by the graph^35,49,53–55^, and often optimize for accuracy at the single-nucleotide level rather than for amenability and interest for analysis. Copangraph instead considers homology between sequences when merging together single-sample sequence graphs (**Fig. 2b**), based on a tunable threshold, providing the user with direct and fine-grained control over the realization of genomic variation in its structure. In this process (**Fig. 2c**), we first detect homologous intervals across the many (potentially millions of) contigs represented by the nodes of each sample-specific sequence graph (**Fig. 2d**). This requires clustering pairwise alignments, which often have variable start- and end-points and different orientations (**Fig. 2e,f**). We developed a clustering algorithm that accomplishes this by clustering the endpoints of each pairwise alignment (**Fig. 2g,h**), first within (**Fig. 2i**) and then across (**Fig. 2j**) contigs, generating “homology groups” (**Methods**).

The final graph is then constructed from these homology groups (**Fig. 2k**). We construct a node for each group, which will represent all the sequence intervals that it includes (**Fig. 2l**). An edge is induced between a pair of nodes if there is a contig or an edge in one of the sample-specific sequence graphs that connects between them (**Fig. 2l**). Copangraph keeps track of the individual sequences that are clustered in each node, as well as the sample of origin for all nodes and edges. This information, made available to the user, enables copangraph to be used as a framework for de-novo comparative analysis of the pan-metagenome. Copangraph is available as an efficient open-source C++ package at https://github.com/korem-lab/copangraph, with a complete technical description provided in the **Methods** section.

### Contiguity detection improves single-sample sequence-graph generation

Since the accuracy of the graph is central to its analytical utility, we quantified how well copangraph represents the underlying metagenome. We started by analyzing the accuracy of copangraph when applied to individual samples, in order to evaluate its single-sample stage (**Fig. 2a**). To this end, we generated ten metagenomes, using CAMISIM^52^ to simulate complexity similar to a gut microbiome (**Methods**). We simulated 80 million read pairs, and then randomly sub-sampled 1-60 million pairs from each sample. We assembled each sample using MEGAHIT^54^, a popular and scalable assembler, and then compared the contigs and assembly graph that it produced with a MetaCarvel^35^ scaffold graph and a single-sample copangraph, with the latter two methods applied to the MEGAHIT contigs (**Methods**).

First, we defined a metric of coverage, which quantifies the number of reference basepairs (bp) that are accurately represented in a sequence graph (**Fig. 3a**; **Methods**). When we quantified the F-score for coverage for each of the different methods, we found that copangraph’s performance was similar to MEGAHIT’s graph and contigs at all depths **(Fig. 3b**). In contrast, MetaCarvel had significantly lower coverage F-score than other methods (Wilcoxon signed-rank *p* = 0.001 vs. copangraph at all depths).

**Figure 3.**
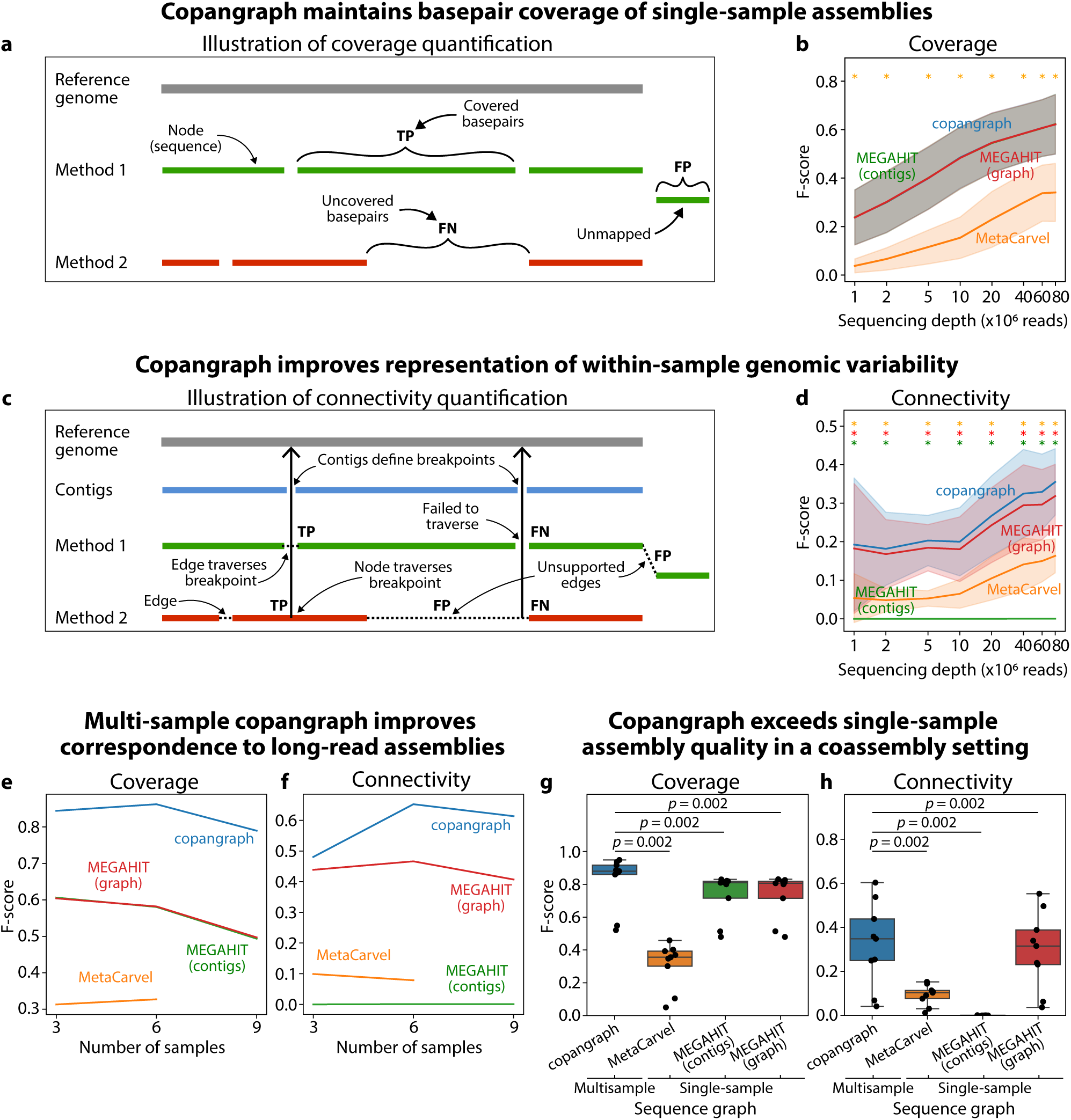
Copangraph accurately represents the sequence and variant information present in metagenomic data. **a,** Illustration of how coverage is quantified. The outputs of various methods are mapped to each ground-truth reference (gray). A reference bp is classified as a true positive (TP) if exactly one sequence maps to it, or as a false negative (FN) if no sequence maps to it. If n > 1 sequences map to the reference, we count one TP and n-1 false positives (FP). Each bp of an unmapped graph sequence is also classified as FP. **b**, The F-score of coverage (y-axis) across a range of sequencing depths (**Methods**). Line, mean F-score across ten simulated samples; shade, ±1 s.d; *, one-sided Wilcoxon signed-rank *p* = 0.001 between copangraph and the method with the corresponding color. **c,** Illustration of how connectivity is quantified. Contrary to coverage, connectivity examines how well a sequence graph represents variability in the underlying genomes. To this end, we examine the genomic positions at the edges of contigs produced by MEGAHIT (i.e., the points in which it terminated assembly paths), as they are most likely to be involved in complex genomic regions such as repeats, variants, or deletions^9,32^ (**Methods**). As in (a), we map genomic sequences output by various methods to each ground-truth reference (gray). Variable genomic positions that are either contained within a single graph sequence or flanked by a pair of sequences that have an edge between them are classified as TP, and, otherwise, as a FN. Edges between sequences that connect distant or unmapped contigs are classified as FP. **d,** same as (b) for connectivity. **e,f,** A benchmark of real human gut metagenomic samples with paired long-read sequencing used as ground truth (**Methods**). F-score of coverage (e) and connectivity (f) of multi-sample copangraph compared to the graph and contigs of MEGAHIT coassemblies, and a MetaCarvel scaffold graph ran on MEGAHIT coassembly. **g**,**h,** The F-score of coverage (g) and connectivity (h) of single-sample subgraphs extracted from a multi-sample copangraph, compared with single-sample assembly graphs and contigs produced by alternative methods. *p*, one-sided Wilcoxon signed-rank test.

Next, we defined a metric of connectivity, which quantifies how accurately a sequence graph represents regions that are repetitive or variable (**Fig. 3c**; **Methods**). We define these regions using contig edges (**Methods**), since assemblers tend to break contigs at these regions^9,32^. Therefore, when we quantified the F-score for connectivity for each of the different methods, we found that, as expected, MEGAHIT contigs have a score of zero across all depths. Copangraph significantly outperformed both MEGAHIT and MetaCarvel graphs across all sequencing depths (Wilcoxon signed-rank *p* = 0.001 for all; **Fig. 3d**). Taken together, these results demonstrate that our approach to single-sample sequence graph generation maintains coverage of the ground truth genomes while significantly improving representation of genomic variability.

### Copangraph accurately represents the pan-metagenome of multiple samples

We next evaluated copangraph in a multi-sample setting, comparing it to coassemblies constructed by alternative methods. We did not use the simulated benchmark in this case, as it does not sufficiently represent strain variability between coassembled samples, which is critical for assembly quality^44^. Instead, we used a dataset of nine gut metagenomic samples from six healthy individuals that underwent both short-read (Illumina NovaSeq 6000), and long-read (PacBio Sequel II) sequencing^56^. In this case, we considered the long reads as a ground truth, and generated low-contamination metagenome-assembled genomes to reduce redundancy and improve quality (**Methods**). We then applied the same methods as above to the short-read metagenomes, building coassemblies of three, six, and nine samples (**Methods**). Evaluating the different methods against the ground truth using our coverage and connectivity metrics (**Methods**), we observed a substantial performance increase for copangraph compared to MEGAHIT coassemblies, MEGAHIT coassembly graphs, and MetaCarvel multi-sample scaffold graphs, for both coverage (**Fig. 3e**) and connectivity (**Fig. 3f**). MetaCarvel, which was included here for completeness despite not being designed for coassembly, did not complete the nine-sample run within our allotted five days. We noted lower coverage and connectivity across methods with an increased number of samples, which is likely explained by increased fragmentation and reduced sequence length, leading to issues with mappability.

Coassemblies have been shown to fall in accuracy compared to single-sample assemblies^43,44^. We therefore next sought to compare multi-sample copangraphs to single-sample assemblies produced by other methods. To test this, we extracted the subgraph and sequences belonging to each sample from the nine-sample copangraph. We then compared their coverage and connectivity F-scores to single-sample MEGAHIT assemblies, MEGAHIT assembly graphs, and MetaCarvel graphs. Copangraph’s subgraphs had significantly higher coverage and connectivity F-score than all alternative methods (Wilcoxon signed-rank *p* = 0.002 for all; **Fig. 3g,h**). Taken together, our results demonstrate that copangraph, applied in a multi-sample, “coassembly” setting, provides a more accurate representation of the underlying metagenomes when compared to alternative methods applied in both single- and coassembly settings.

### Copangraph structure reflects phylogeny

After establishing the accuracy of copangraph, we next sought to evaluate how well its structure represents phylogeny. A key factor determining copangraph’s structure is the homology threshold (**Fig. 2**). We therefore first evaluated how this threshold affects the resulting structure. Microbial genomic variability manifests through a variety of mechanisms, resulting in sequence divergence that could vary substantially across a genome (e.g., when comparing two recently divergent genomes, conserved genes may have 100% identity, while a horizontal gene transfer could result in 0% identity). To perform an analysis in which sequence divergence is applied uniformly along the genome, we used a simulation under a neutral evolutionary model. We used BacMeta^57^ to simulate populations of 25 bacterial genomes with different mean average nucleotide identity (ANI) among them, creating ten independent replicates for each ANI value (**Methods**). For each 25-member population, we constructed multiple copangraphs with a range of homology thresholds, and counted the number of nodes created by copangraph. We found that copangraph’s representation of each population underwent sharp transitions when its homology threshold matched the average ANI: as expected, populations with a mean ANI below the homology threshold were represented by 25 nodes, each containing one genome. In contrast, those with a mean ANI above the threshold were represented by a single node containing all 25 genomes (**Fig. 4a**).

**Figure 4.**
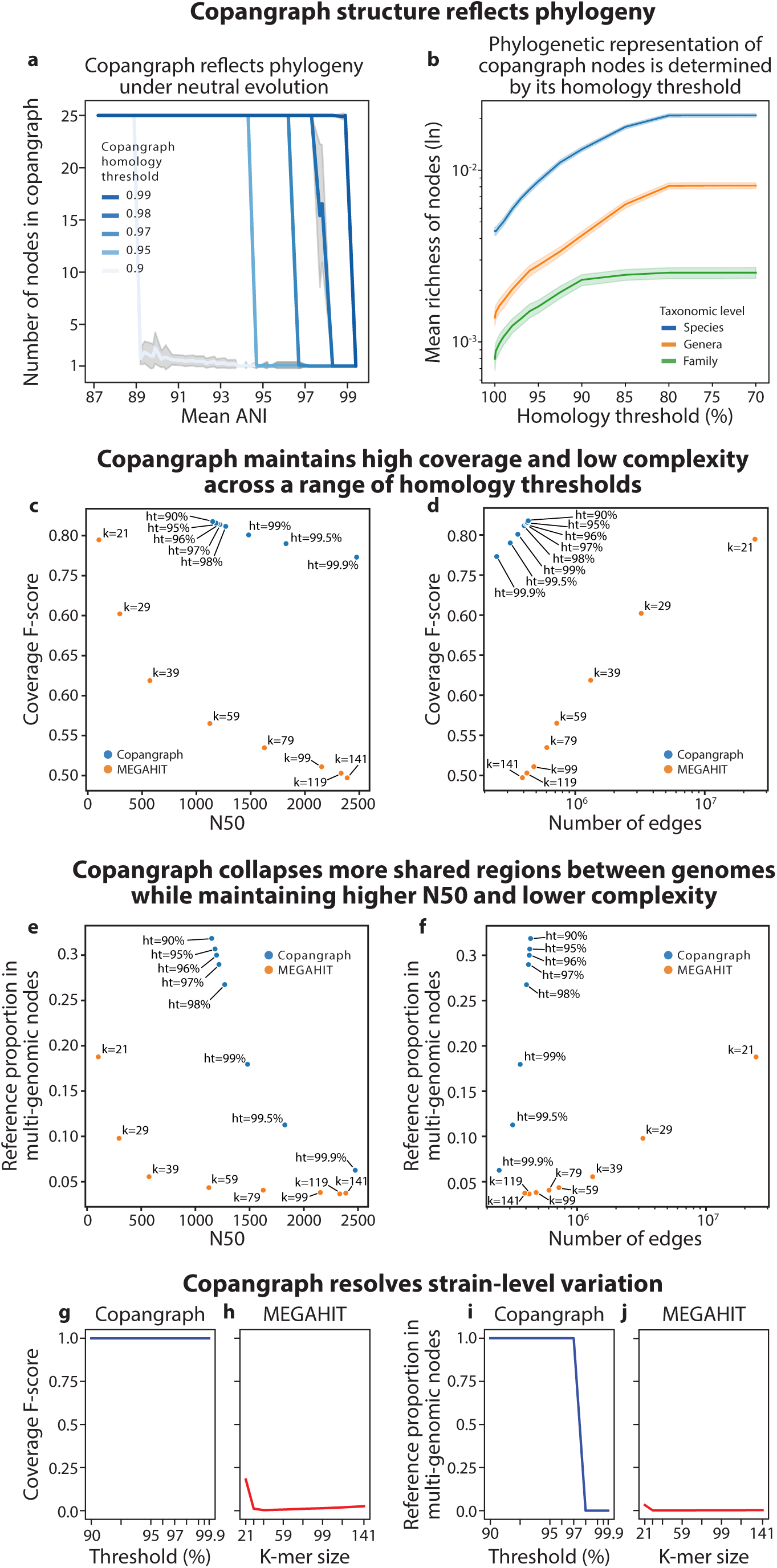
Copangraphs are more suitable for comparative analysis than de Bruijn graph coassemblies. **a**, The number of nodes (y-axis) in copangraphs with different homology thresholds for populations of 25 genomes simulated under a neutral evolutionary model for a specified level of mean ANI (x-axis). When copangraph’s homology threshold matches the population mean ANI, a transition is visible between one node (all genomes collapse) and 25 nodes (every genome is represented by a separate node). Line, mean of ten replicates; shade, 1 s.d. **b**, Mean richness of nodes (y-axis) at different taxonomic levels for copangraphs of ten gut metagenomes simulated with CAMISIM (**Methods**) across different homology thresholds (x-axis). Line, mean; shade, 1 s.e.m. **c**,**d**, Coverage F-score (y-axis; **Methods**) versus N50 (c) and edge count (d) for copangraph and MEGAHIT^54^ coassemblies of nine metagenomic samples from a dataset of paired short- and long-read gut metagenomes^56^. Each point represents a graph, with k- mer size and homology threshold labeled for MEGAHIT and copangraph, respectively. **e**,**f**, Same as c,d (respectively), but showing the proportion of the ground-truth reference metagenome which is present in nodes that represent sequences from multiple genomes. **g,h**, Coverage F-score for copangraph (g) and MEGAHIT coassembly graphs (h), as a function of homology threshold and k-mer size, respectively, for a simulation of 25 genomes at an average ANI of 97%. Line, mean across three replicates; shade, 1 s.d. **i,j**, Same as g, h, respectively, but showing the proportion of reference in multi-genome nodes.

We next evaluated the relationship between the homology threshold and phylogenetic representation in more realistic settings. As our CAMISIM simulations (**Fig. 3**; **Methods**) contained ground-truths with complete taxonomy while reflecting the complexity of the gut microbiome, we used our ten simulated samples to construct multi-sample copangraphs using a range of homology thresholds. We computed the species, genera, and family richness for each node of each copangraph (**Methods**) and found that as we reduced the homology threshold, the richness of the nodes increased across all taxonomic levels (**Fig. 4b**), indicating that more sequences from different clades were collapsed into common nodes. These results demonstrate that the structure of copangraph reflects phylogenetic relationships among the input community, and can be effectively tuned by altering its homology threshold.

### Copangraphs are more suitable for comparative analysis than de Bruijn graphs

The suitability of a sequence graph for comparative analysis depends on multiple factors, with an ideal graph having high coverage, so that it adequately reflects the underlying metagenomes; low fragmentation, meaning that longer sequences are represented by each node, increasing interpretability; lower complexity, reducing computational burden and facilitating analyses; and a high tendency to collapse shared regions across different genomes, which improves detection of both conserved and variable regions^9^. There are often tradeoffs between these factors; for example, achieving higher coverage can be expected to come at the price of higher complexity, and collapsing more shared genomic regions would tend to increase fragmentation. Most sequence graphs therefore have key parameters, such as the k-mer size for de Bruijn graphs, which enable one to tune these graph properties for a specific analysis. We next evaluated these key graph properties for copangraph and MEGAHIT^54^ as a function of homology threshold and k-mer size, respectively, using the coassemblies of nine samples from the dataset of paired short- and long-read metagenomes analyzed above^56^. We evaluated coverage as before (**Fig. 3a**), fragmentation using N50, complexity using the number of edges, and the tendency of the graph to collapse shared regions by calculating the proportion of the ground-truth reference metagenome that is present in nodes that represent sequences from multiple genomes (**Methods**).

First, examining coverage, we found that as k-mer size and homology threshold increased, coverage and graph complexity decreased while N50 increased (**Fig. 4c,d**). This was particularly prominent for MEGAHIT: as k-mer size increased, we observed an order-of-magnitude increase in N50 (103 to 2,388 bp; **Fig. 4c**) and a two-orders-of-magnitude decrease in the number of edges (2.4×10^7^ to 3.9×10^5^; **Fig. 4d**). However, the increased N50 and reduced graph complexity came with a substantial drop in coverage F-score, from 0.79 at k=21 to 0.50 at k=141. While copangraph followed the same general trends, its characteristics varied in a more favorable range: 0.77 to 0.82 coverage (higher than all MEGAHIT graphs except for k=21), 1,150 to 2,475 bp for N50, and 2.5×10^5^ to 4.3×10^5^ edges (**Fig. 4c,d**). Overall, we found copangraph maintained high coverage and low complexity across a broad range of homology thresholds.

Next, examining the proportion of reference genomes in multi-genome nodes, we found similar trends of negative correlation with k-mer size and homology threshold (**Fig. 4e,f**). While in this case copangraph spanned a range more similar to MEGAHIT (0.06-0.32 for copangraph vs. 0.04-0.19 for MEGAHIT), most copangraph homology thresholds led to high proportion in multi-genome nodes (0.27-0.32 for copangraph with homology threshold ≤98%), while most k-mer sizes led to low proportions (0.04-0.06 for MEGAHIT with k≥39). Furthermore, we found that copangraph could represent a similar proportion in multi-genome nodes as MEGAHIT at a substantially larger N50 and lower graph complexity (**Fig 4e,f**). For example, a copangraph with a homology threshold of 99% had a proportion of 0.18 in multi-genome nodes, with N50 of 1,481 bp and 3.6×10^5^ edges; achieving a similar proportion of 0.19 with MEGAHIT, however, required setting k=21, which yielded an N50 of 103 bp and 2.4×10^7^ edges. Taken together, copangraphs demonstrated a consistently more desirable sequence-graph property regime than de Bruijn graphs across a broad range of k-mer sizes and homology thresholds.

### Copangraphs provide a better representation of similar strains than de Bruijn graphs

Strain-level diversity often drives host-microbiome interactions^11–13,15,58^ while at the same time reducing assembly quality and increasing its fragmentation^43,44^. This phenomenon could be exacerbated in a coassembly setting, in which strains from different samples are pooled together. We therefore next sought to directly evaluate copangraph’s ability to represent strain-level variation, using three replicates of 25-genome simulations used above with a mean ANI of 97%. To model cross-sample strain heterogeneity, we treated each genome as a separate sample, simulated paired-end reads from it, constructed multi-sample copangraphs, and quantified the coverage F-score and proportion of reference genome in multi-genome nodes. We found that copangraph provided perfect coverage of each strain regardless of the homology threshold used (**Fig. 4g**). In contrast, MEGAHIT coassembly graphs showed low coverage F-score, with a maximum of 0.18 at k=21, and minimum of 0.003 at k=39 (**Fig. 4h**). Examining the proportion of reference in multi-genomic nodes, we found that copangraph appropriately collapsed all genomes to a single node when its homology threshold was set to 97% or less (**Fig. 4i**). In contrast, MEGAHIT provided substantially lower collapsing across genomes, with a range of 8.6×10^-5^ to 0.032 (**Fig. 4j**). Overall, this analysis demonstrates that copangraph is able to appropriately represent strain-level variation under a neutral evolutionary model.

### Copangraph construction scales efficiently with increasing samples

We next wished to profile the computational requirements of copangraph construction. To this end, we sequenced gut metagenomic samples from a liver transplant cohort^14,50^ (**Methods**) and built copangraphs of random subsets of 5-100 samples. For comparison, we constructed MEGAHIT^54^ and metaSPAdes^55^ coassemblies on the same subsets. We ran each construction in triplicate and recorded runtime and peak memory usage (maximum resident set size; **Methods**). Due to prohibitively long runtimes, we limited the maximum runtime of metaSPAdes to 12 hours, the maximum runtime recorded for MEGAHIT. With this time constraint, metaSPAdes could complete two of the three 20-sample coassemblies, but not any larger coassemblies. Copangraph construction was faster and had lower peak memory usage than MEGAHIT coassembly for all replicates with 20 or more samples and all metaSPAdes runs (**Fig. 5**). While copangraph scaled slightly better than MEGAHIT, both appeared to scale linearly with the number of samples (**Fig. 5**). These results demonstrate the scalability of copangraph construction, showing that it is slightly faster than even a highly efficient de Bruijn graph assembler.

**Figure 5.**
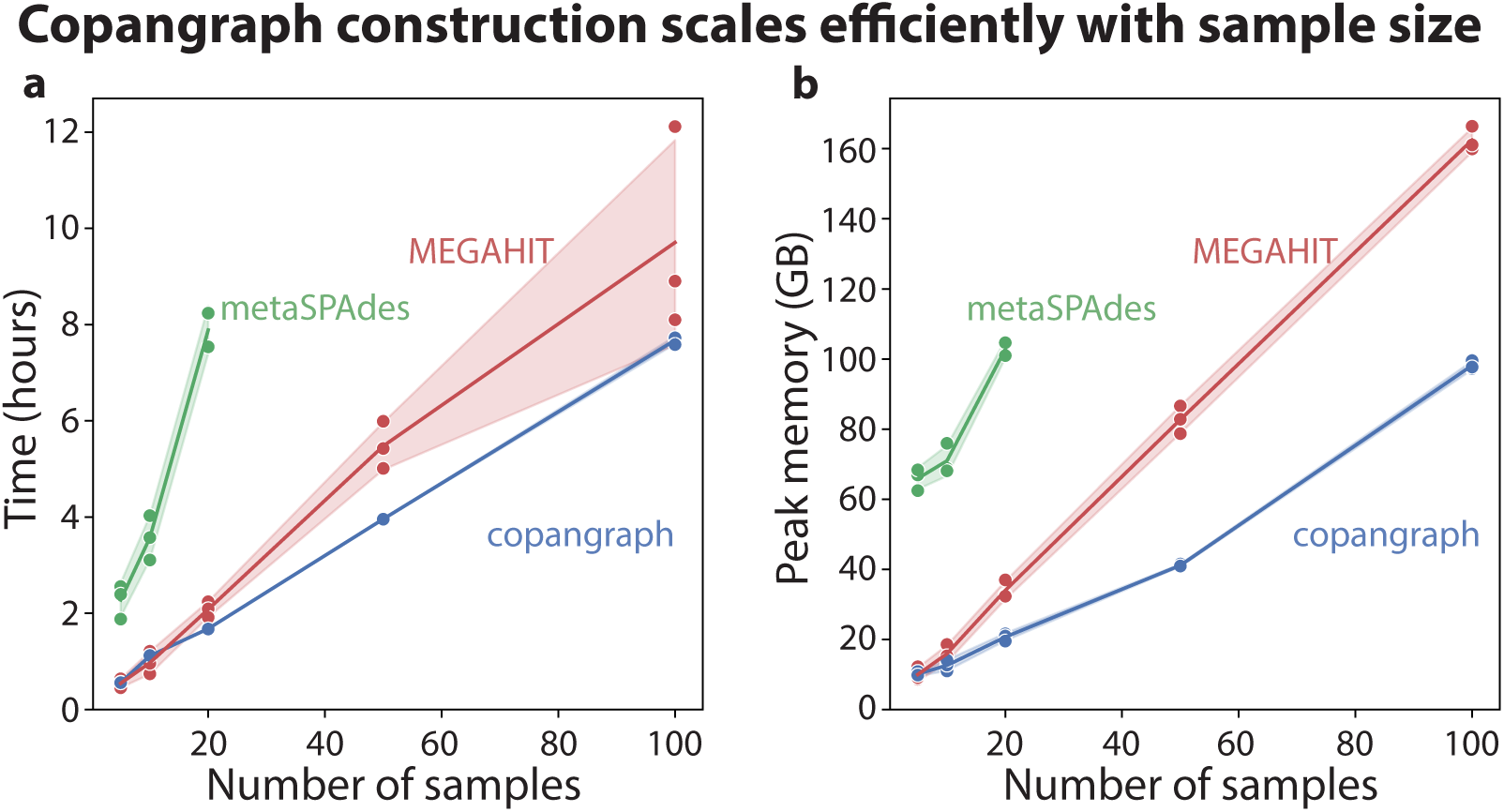
Copangraph construction scales to large sample sizes. **a,b**, The runtime (a) and peak memory usage (resident set size) (b) of copangraph construction compared to MEGAHIT and metaSPAdes coassembly construction at increasing input sample sizes (line, mean; shade, 1 s.d.) MetaSPAdes runtime was capped at 12 hours.

### Copangraph provides an informative microbiome representation in diverse settings

We next assessed the utility of copangraph in representing variability across metagenomes across different settings. To this end, we obtained gut metagenomic data collected from patients and controls in a study of colorectal cancer^59^ (N=128; 75 with colorectal cancer) as well as vaginal metagenomic data collected during the second trimester of pregnancies (16-24 weeks of gestation) that ended preterm spontaneously and at term^60^ (N=100; 29 delivered preterm; **Methods**). We constructed a multi-sample copangraph from the metagenomic data of each, and calculated Jaccard and Euclidean distances over its variable regions (**Methods**). We compared how much of the variability in these distances was explained by each phenotype, and compared them to Bray-Curtis dissimilarities calculated over mOTUs3 (ref. ^61^) species abundances, a common practice in the field. We found that 4.7% of the variance in Jaccard distances calculated from a gut metagenomic copangraph was explained by the presence of colorectal cancer (PERMANOVA p<0.001), as opposed to 1.9% of the variance in Bray-Curtis dissimilarities calculated over mOTUs3 abundances (**Fig. 6a,b**), while only 1.7% of the variance in copangraph Euclidean distances (p=0.12; **Fig. 6c**). Examining spontaneous preterm birth, we found that once again it was more strongly associated with Jaccard distances calculated from a vaginal metagenomic copangraph than with Bray-Curtis dissimilarities calculated from mOTUs3 abundances (3.5% vs. 2.6%; p=0.016 and 0.014, respectively; **Fig. 6d,e**). In this case, however, 9.1% of the variance in copangraph Euclidean distances was explained by spontaneous preterm birth (p=0.002; **Fig. 6f**).

**Figure 6.**
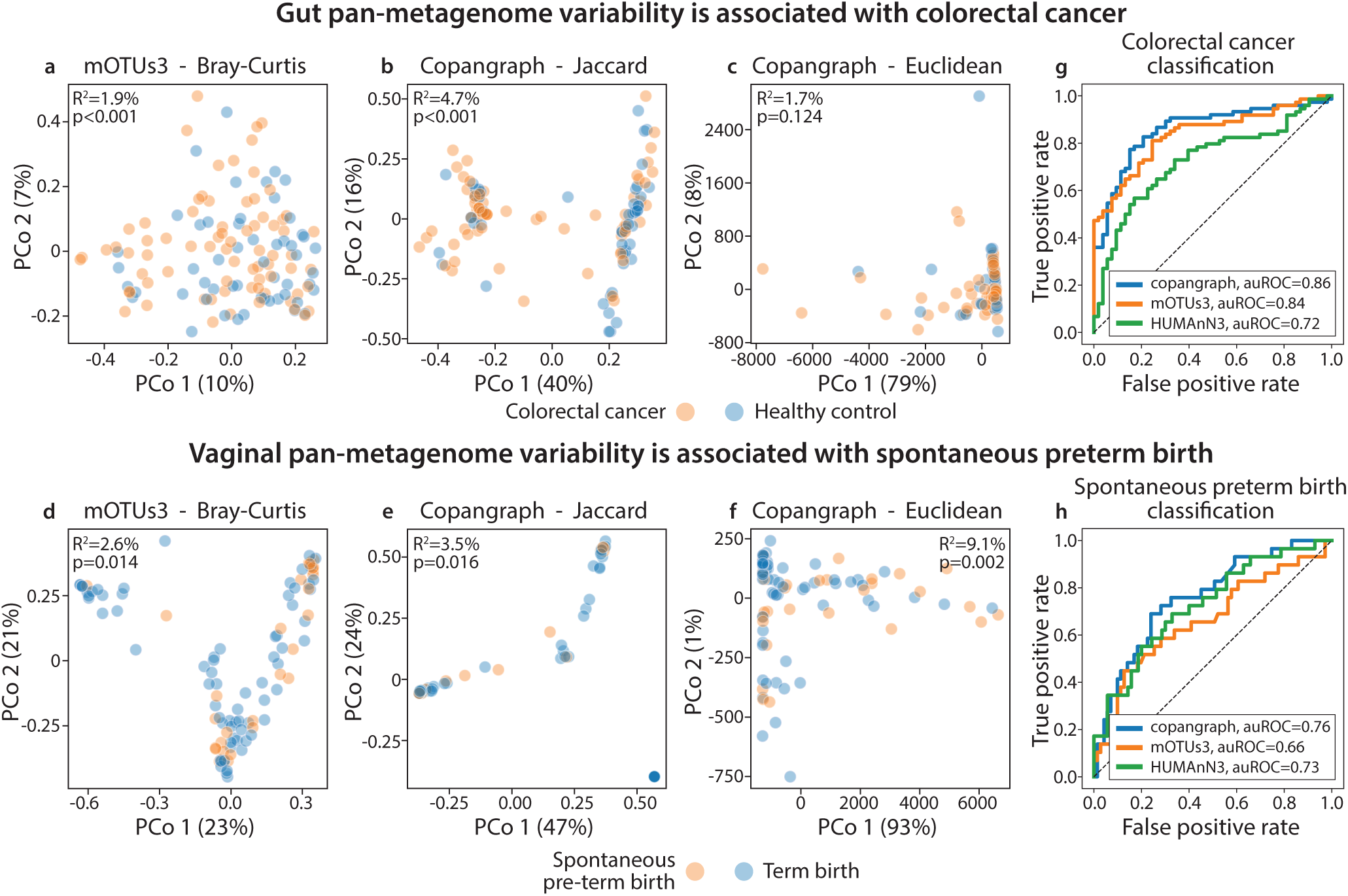
Copangraph represents metagenomic variability across different scenarios. **a-c**, For an analysis of gut metagenomic samples from a colorectal cancer study^59^, shown are the first two Principal Coordinates (PCo) from a Principal Coordinate Analysis (PCoA) using Bray-Curtis dissimilarity calculated using species abundances (a), Jaccard distances calculated from copangraphs (b), or Euclidean distances calculated from copangraph (c). ADONIS^65^ explained variance and p value is displayed on each panel. **d-f,** Same as **a-c** for vaginal metagenomic samples from a study of spontaneous preterm birth^60^. **g,** Receiver operating characteristic (ROC) curves for classification of colorectal cancer, comparing similar classifiers based on copangraph (auROC=0.86), mOTUs3 (ref. ^61^, auROC=0.84) and HUMAnN3 (ref. ^62^, auROC=0.72). **h**, ROC curves for classification of spontaneous preterm birth, comparing similar classifiers based on copangraph (auROC=0.76), mOTUs3 (auROC=0.66) and HUMAnN3 (auROC=0.73).

To further evaluate the information content of copangraph-based representation, we performed a prediction benchmark, comparing a tabular representation of copangraphs to mOTUs3 (ref. ^61^) species abundances and HUMAnN3 (ref. ^62^) gene abundances. For all three feature types we trained and evaluated a gradient-boosted decision-trees classifier using a nested 10-fold cross-validation (**Methods**). Evaluating classification of colorectal cancer, we found high predictive performance using mOTUs3 (area under the receiver operating characteristic curve [auROC] of 0.84), in line with previous reports^63,64^, and somewhat weaker performance for prediction based on HUMAnN3 (auROC=0.72) (**Fig. 6g**). Copangraph-based prediction was similar and slightly higher, with an auROC of 0.86 (**Fig. 6g**). When evaluating classification of spontaneous preterm birth HUMAnN3 gene abundances had higher performance than mOTUs3 species abundances (auROC of 0.73 vs. 0.66, **Fig 6h**). Here also, copangraph based prediction was similar and slightly better, with an auROC of 0.76 (**Fig. 6h**). Overall, this analysis demonstrates that copangraphs is consistently able to capture and represent phenotypically meaningful metagenomic variability across different ecosystems and clinical scenarios.

### Copangraph enables prediction of VRE colonization trajectory

Multi-drug resistant (MDR) colonization of the gut is very prevalent in liver transplant recipients^14^. While it significantly increases the risk of transmission^14^ and subsequent infections^14,50,66,67^, most patients will clear colonization spontaneously^14^. Identifying patients at high risk for colonization persistence and infection would facilitate prophylactic treatment. Having designed copangraph for comparative analysis of samples in a pan-metagenome (comparative metagenomics), we next sought to assess its effectiveness at predicting the colonization trajectory of Vancomycin-resistant *Enterococcus* (VRE), an MDR ESKAPE pathogen^68^ (**Fig. 7a**). To this end, we performed metagenomic sequencing of 77 stool samples from 62 liver transplant recipients who participated in a prospective cohort with longitudinal follow-up^14,50^ (**Table S1**). Each sample was previously confirmed to be VRE positive using selective culture (**Methods**). An additional VRE culture result was obtained between three days to four weeks after the first sample was taken, which we used to define VRE colonization trajectory: persistence if the sample was culture positive, or clearance if it was negative. We constructed a copangraph from this metagenomic data collected alongside the first VRE-positive culture (**Fig. 7a**). Examining overall association of copangraph representation and colonization trajectory, we found a statistically significant association with Jaccard distances calculated from copangraphs (2.9% explained variance, PERMANOVA p=0.017), and no significant associations with mOTUs3 Bray-Curtis dissimilarities or copangraph Euclidean distances (1.1%, p=0.69 and 1.2%, p=0.64, respectively; **Fig. S1**).

**Figure 7.**
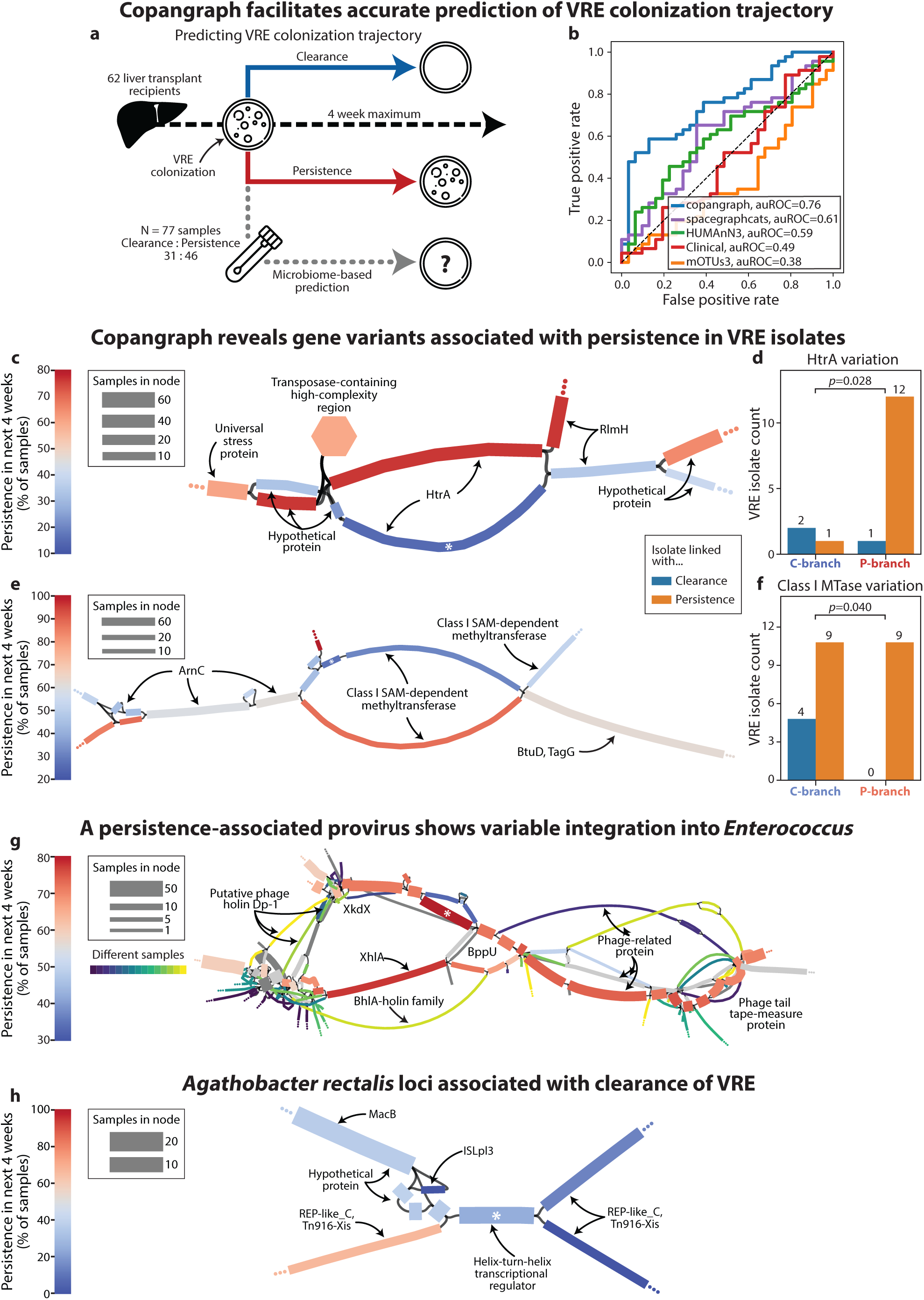
Comparative metagenomics of VRE colonization trajectory using copangraph. **a**, Schematic of our analysis of Vancomycin-resistant *Enterococcus* (VRE) colonization trajectory. Our cohort includes 77 independent timepoints in which 62 liver transplant recipients had culture-confirmed gut VRE colonization. Using metagenomic sequencing data from the same time, we attempt to predict whether colonization would clear spontaneously (N=31) or if it would persist (N=46), determined using another culture within the subsequent four weeks (**Methods**). **b**, Receiver operating characteristic (ROC) curves for prediction of VRE colonization trajectory, comparing similar classifiers based on copangraph (auROC=0.76), mOTUs3 (ref. ^61^, auROC=0.38), HUMAnN3 (ref. ^62^, auROC=0.59), spacegraphcats^53^ (auROC=0.61), and clinical (auROC=0.48) features. **c**, Copangraph subgraph surrounding a predictive node that is part of the *htrA* gene in VRE. **d**, Barplot tallying the number of VRE isolates to which each branch of the *htrA*-containing bubble in (c) mapped at ≥ 99.5% identity, stratified by isolates from samples followed by VRE persistence and clearance. “C-branch” and “P-branch” refer to the two *htrA* branches in copangraph with a higher proportion of samples followed by clearance and persistence, respectively. *p*, one-sided Barnard’s exact. Neither branch mapped at ≥99.5% identity to seven of the 23 isolates. **e**, Copangraph subgraph surrounding a predictive node, part of a gene coding for a class I SAM-dependent methyltransferase. **f**, Same as d, for the two branches of the methyltransferase-containing bubble in (e). Neither branch mapped at ≥99.5% identity to one isolate. **g**, Copangraph subgraph surrounding a predictive node that is part of an Enterococcal prophage element. To highlight the prophage, only nodes annotated as prophage (**Methods**) display the percentage of samples followed by VRE persistence. To demonstrate the variable integration of this phage across samples, nodes containing sequences from just a single sample were given a color unique to that sample. **h**, Copangraph subgraph surrounding a predictive repeat region in *Agathobacter rectalis*. All sequences in each node of c,e,g blasted to *E. faecium* or *E. faecalis* with sequence identity > 99.5%. For h, the feature and *macB* nodes blasted to *A. rectalis* with 100% identity, the ISLpl3 node to *Ruminococcus torques* at 100% identity, and the *Tn916* containing nodes did not map to any genome in ncbi nr^71^ with identity above 92% (the best matches were to *Blautia coccoides*). For c,e,g,h: node color, percentage of samples in node that were followed by VRE persistence; node thickness, total number of samples in node; ellipsis, subgraph extends to the rest of the graph; asterisk, predictive node.

Next, we trained and evaluated a gradient-boosted decision-trees classifier using nested 10-fold cross-validation (**Methods**). For comparison, we trained similar classifiers using mOTUs3 (ref. ^61^) species abundances, HUMAnN3 (ref. ^62^) gene abundances, all available clinical covariates^50^, and “graph piece abundances” from spacegraphcats^53^ (**Methods**). Prediction of colonization trajectory is a challenging task, and classifiers based on mOTUs3, HUMAnN3, clinical and spacegraphcats features all performed poorly (auROCs of 0.38, 0.59, 0.49, and 0.61, respectively; **Fig. 7b**). In contrast, the copangraph-based classifier performed substantially better (auROC=0.76). These results demonstrate the strong potential of the microbiome for predicting colonization trajectories, highlight the importance of feature engineering, and show that copangraph provides a highly informative representation of the pan-metagenome that aids in difficult prediction tasks.

### Copangraph-based comparative metagenomics reveals specific genomic regions associated with VRE colonization trajectory

We next examined whether our prediction scheme reveals specific genomic regions with potential importance for VRE colonization, facilitating comparative metagenomics. To this end, we quantified the feature importance of nodes and edges in our models and extracted and annotated the subgraphs around the top five features (**Methods**). One such feature was part of the *htrA* gene within *Enterococcus faecium*. Copangraph detected two alternative versions of this gene, manifesting as two distinct branches that formed a “bubble”^69^ in copangraph (**Fig. 7c**). One branch displayed a significantly higher percentage of samples with subsequent persistent colonization than the other (76% vs. 15%; Barnard exact *p*=2×10^-6^). *htrA* was recently found to be involved in endocarditis and biofilm-associated pilus biogenesis, and to contribute to persistent colonization of *E. faecalis*^70^. To validate that the variation detected by copangraph reflects VRE strain variation, we sequenced 23 isolates cultured from the same samples, and performed long-read nanopore sequencing followed by assembly (**Methods**). Mapping one sequence from each branch to these VRE isolate genomes, we found that each mapped to a different and disjoint set of isolates, with the persistence-associated branch mapping to a significantly higher number of persistence-associated VRE strains (*p*=0.028; **Fig. 7d**).

Another top feature was part of a gene containing a class I SAM-dependent methyltransferase domain within *E. faecium*. Once again, copangraph detected two versions of this gene, which formed a bubble, with one branch having a significantly higher percentage of samples with subsequent persistent colonization (88% vs 28%; Barnard exact *p*=2×10^-6^; **Fig. 7e**). To validate this structure experimentally, we performed the same analysis of VRE isolates. Once again, we found that each branch mapped to a different and disjoint set of isolates, with a significant association with the subsequent colonization trajectory of the VRE isolates (*p*=0.040; **Fig. 7f**). Overall, this analysis demonstrates that copangraph can detect biologically relevant genetic variation in VRE strains that is associated with subsequent colonization trajectory.

Other top features used by the copangraph-based classifier were not specific to VRE. We found one top feature which was part of a prophage dispersed across a chain of copangraph nodes, all of which were present predominantly in samples that were followed by subsequent VRE persistence (**Fig. 7g**). The “core” prophage chain, defined as nodes annotated as a prophage by geNOMAD^72^, was surrounded by additional nodes, many of which contained phage-related genes (but not annotated as prophage elements). The prophage and surrounding subgraph were embedded in a region of the graph annotated as *E. faecium* (**Methods**), with substantial cross-sample variability, suggesting that this prophage is integrated variably within this species. Even the prophage itself showed a variable structure, containing two major branches that were both annotated as part of the prophage, and both had a higher proportion of samples followed by persistence. One of these two branches harbored an *xhlA* gene, which was previously shown to form part of a lytic cassette with variable presence in Enterococcal phages^73^. Interestingly, the predictive node itself was part of a structural variation (seen as a bubble), with the feature branch having a significantly higher proportion of samples with subsequent persistence than the other (83% vs 33%; Barnard exact *p*=0.003).

Finally, copangraph identified a genomic region in *Agathobacter rectalis*, which we found predominantly in samples with subsequent clearance (75%; Barnard exact *p*=0.0015; **Fig. 7h**). The feature was part of a helix-turn-helix transcriptional regulator, and was flanked by the *Tn916* transposon genes *REP-like_C* and excisionase *Tn916-Xis*, as well as by *macB*, an ABC transporter conferring antibiotic resistance^74^. *A. rectalis*, formerly *Eubacterium rectalis*, is a known butyrate producer^75,76^ associated with gut health^77^. These results demonstrate the importance of examining the surrounding microbiome that a VRE strain colonizes, which may affect colonization trajectory and potentially confer some resistance, echoing our previous results that found microbiome associations with colonization resistance^50^.

## Discussion

To bridge the gap between comparative genomics and microbiome studies, we extended sequence-graph analysis to the comparative metagenomic setting by introducing copangraph (comparative pan-metagenomic graph). Copangraphs are multi-sample homology-based sequence graphs constructed by hybrid coassembly, a novel approach that constructs accurate single-sample graphs and then efficiently merges them together. We introduced several novel algorithms for single-sample sequence-graph reconstruction, merging, and collapsing of homologous regions. We demonstrated that copangraphs offer a faithful multi-sample representation of the underlying pan-metagenome, with accuracy surpassing sample-specific representations. We showed that copangraph construction is tractable and scalable, and provides a multi-sample sequence-graph that is more amenable for comparative analysis than de Bruijn graphs. Copangraph demonstrated the ability to consistently capture phenotypic associations in diverse clinical scenarios. Particularly in the challenging clinical and biological scenario of predicting VRE colonization trajectory in liver transplant recipients, copangraph provided a more informative representation of the microbiome than species or functional abundances, which enabled accurate prediction. Finally, we performed a comparative metagenomic analysis, identifying specific genomic regions that are strongly associated with subsequent VRE persistence and clearance, with significant association with isolated VRE strains. Overall, copangraph offers a new and useful conceptual and practical framework for *de novo* pan-metagenomic analyses.

The diversity of microbial communities and the complex variability of their genomes warrant the development of sufficiently expressive and reference-free comparative frameworks. When combined with scalable approaches for construction and analysis, sequence graphs are well-suited for such a framework; therefore, recent efforts have developed sequence-graph-based methods for comparative analysis^45,47,53^. However, challenges with scalability and complexity have so far set the focus on comparing single-sample graphs from different environments^45,47^, or on coarse-graining of multi-sample graphs^53^. The use of homology-based graphs allows us to collapse evolutionary-related sequences together in the same node, providing a biologically motivated, rather than arbitrary, coarse graining.

This is further facilitated by hybrid coassembly, a new approach for *de novo* metagenomic analysis that combines the accuracy and scalability of single-sample assembly with the capacity for comparative analysis afforded by coassembly. The separation of assembly from multi-sample graph construction by hybrid coassembly offers substantial flexibility for de-novo analysis, allowing the study of genomic variability at different scales, from variation at the SNP level to larger-scale genomic rearrangements^78^. Such variation need not necessarily even be defined based on DNA sequence homology, and future work could explore graph merging based on protein sequence homology or language-model embeddings^79^, which may result in comparative multi-sample graphs with different analytical advantages. While copangraph is a specific implementation of hybrid coassembly, the approach is generalizable and could be applied for de Bruijn graph-based merging; long-read metagenomic samples, which may offer comparative advantages as well as framework for integration with short-reads; or even metatranscriptomics^80^.

Taxonomic and gene profilers are efficient and provide informative metagenomic representations; recent developments also facilitate their use in the context of *de novo* assembly^61,81^. However, our results indicate that even in very well-studied ecosystems such as the human gut, profilers can produce less informative representations than pan-metagenomic representations for some phenotypes. While in some scenarios we tested species or functional abundances performed comparably to copangraph, ours was the only representation that both performed well consistently and facilitates comparative analysis of variable genomic regions. Our results underscore the challenges in translating genomic variability to coherent taxonomic definitions^82^, and the value in data-driven, sequence-graph-based approaches.

A notable such approach is spacegraphcats^53^, which performs coassembly on a combination of sample-specific read-sets that are enriched to the sample-specific “neighborhoods” of taxa of interest, which are queried from sample-specific coarse-grained assembly graphs. To handle the complexity of this task, coarse-graining is applied at multiple stages in the form of r-dominating sets^48,53^. The difference we observe in the performance of spacegraphcats and copangraph is likely explained by the different approaches used for coarse-graining (r-dominating sets vs. homology, respectively), coassembly (de Bruijn graph-based coassembly vs. hybrid coassembly), and the focus of spacegraphcats on specific taxa.

In comparative analyses, the most appropriate taxonomic or genomic resolution for analysis is impacted by data modality, data quantity, and the particular phenomena under study^78,82–84^. This is evident, for example, in prediction, whose performance can be substantially impacted by the taxonomic resolution of the data used^84,85^. The optimal genomic resolution for analysis could be impacted by the evolutionary rate of closely-related genomes, which is affected by environmental conditions^86^, as well as by phenomena that can substantially alter genome structure, such as horizontal gene transfer^39^. We find it likely that an optimal sequence identity for analysis does not exist, and therefore, comparative frameworks should facilitate analysis at different resolutions. In de Bruijn graphs, altering the k-mer size impacts the tendency for shared regions to collapse, but, as we show, it is also strongly related to graph complexity, sequence length, and coverage. In copangraph, the tunable homology threshold is largely decoupled from these properties, enabling comparative analyses of the pan-metagenome at different resolutions without dramatic shifts in complexity, sequence length, or coverage. We anticipate that this tunability will enable copangraph to be an informative representation across a broad range of microbiome-related analyses. Still, the current version of copangraph enables using only a single constant threshold per graph, requiring potential tuning. An adaptive threshold that respects sequence or local graph topology may better account for variability in ANI across taxa or genomic regions, and is a promising future research direction.

We have demonstrated the utility of copangraph in a comparative metagenomics analysis of a challenging clinical scenario. We note, however, that the representation of nodes and edges in our predictive analysis was a simple tabular representation, and that copangraphs could serve as the basis for other approaches, such as graph embedding methods^87^. Alternatively, an analysis driven by the identification of known topological signatures of genomic events, such as bubbles^88^, can drive a broader investigation of the genetic events they represent. Furthermore, given the high accuracy of copangraph in representing the pan-metagenome, it could be useful for a variety of other downstream analyses, such as for calculating sensitive similarity metrics between microbiomes, as the basis for graph-informed binning approaches^89^, or for strain-level analysis methods that combine the syntenic information coded in the graph with the nucleotide-level variability present within its nodes. To facilitate these further uses of copangraph, we have created an efficient C++ implementation of copangraph available at https://github.com/korem-lab/copangraph, with detailed usage instructions.Altogether, copangraph offers a versatile framework for comparative pan-metagenomics that facilitates high-resolution microbiome investigations.

## Methods

### Copangraph construction

#### Single-sample stage

Copangraph construction begins with a single-sample stage that constructs a sequence graph for each sample (**Supplementary Algorithm 1**: `buildSampleSequenceGraph`). First, we assemble each short-read paired-end metagenomic sample separately. In this work, we use MEGAHIT^54^, which is fast and accurate, but note that other assemblers^55,90–92^ could also be used. Specifically, we used MEGAHIT v1.2.9 in default settings, and used the final.contigs.fa file. We then construct a sequence graph for each sample by detecting pairs of contigs that represent adjacent genomic regions. We start by assembling the genomic regions immediately up- or downstream of each contig, using a local assembly algorithm that makes full use of paired-end information (**Supplementary Algorithm 2**: `extendContig`). This algorithm requires a mapping of each sample to its contigs, which we obtained using Bowtie2 v2.5.3 (ref. ^93^) in paired-end mode to construct a .bam file, which we subsequently sorted by read name using samtools v1.18. For each extension assembled for a particular contig, we then search for high-confidence prefix-suffix overlaps with any other contig in the assembly. For each detected overlap, we add a bi-directed edge^94^ between the two contigs to the sequence graph, which specifies the complement (forward or reverse) of the two contigs in the detected overlap.

#### Tunable graph merging: creating homology groups

After a sequence graph was created for each sample, we merge all of these graphs together into a multi-sample copangraph. We do this by identifying homologous genomic regions (i.e., intervals within contigs / nodes) across the graphs and collapsing them into new, multi-sequence-multi-sample nodes. We detect these cross-sample homologies using pairwise alignments between the contigs of all samples, which poses several challenges: (1) a genomic region may have multiple different alignments that overlap, but have different start and end points; (2) homologous intervals can be in opposite complement to one another; (3) a larger homologous genomic region can be composed of smaller regions that are themselves homologous to different contigs, forming complex “mosaic repeats” (**Figs. S2,3**); and (4) homology (over some cutoff; and similarly for pairwise alignment) is not transitive, meaning that region A may be homologous to region B, and B to C, but A and C would not be homologous. We term the groups of cross-sample homologous genomic intervals “homology groups”, and represent each one with a node. We designed homology groups to meet a few criteria: (a) the intervals in each group are homologous, that is, share a sequence identity above a user-specified threshold with at least one other interval in the group; (b) the intervals are disjoint from all other intervals in the graph, meaning that different intervals never overlap; and (c) the intervals provide complete coverage, meaning that all intervals in the input contigs are represented by a homology group.

To construct homology groups achieving these criteria (**Supplementary Algorithm 3:** `createHomologyGroups`), we first compute pairwise alignments between the contigs, using an efficient, minimizer-based co-linear chaining algorithm inspired by minimap2 (ref. ^95^). The pairwise alignments are then filtered, discarding those with sequence identity below the user-specified homology threshold and minimum alignment length. Next, we cluster these pairwise alignments into homology groups. However, because intervals may have different and overlapping start and end points, we do not cluster the intervals directly, but instead cluster their endpoints, arriving at consensus disjoint start and end positions of all the intervals that would divide each contig.

We start by turning each alignment into four endpoints, with each endpoint specifying a position on a contig and the relative position of the aligned interval with respect to the endpoint - whether the interval is to the endpoint’s *left* (toward position 0 of the contig) or to its *right* (toward the end of the contig). Let *s*[*i*: *j*] and *t*[*k*: *l*] be a pair of homologous intervals. We generate two endpoints for *s*: *s*_1_ = (*i, right*) and *s*_2_ = (*j, left*), and two for *t*: *t*_1_ = (*k, right*) and *t*_2_ = (*l, left*). If *s* and *t* align to each other in the same DNA complement, we maintain a “link” between *s*_1_ and *t*_1_, signifying that they originate from the beginning of the same alignment, and likewise for *s*_2_ and *t*_2_, which are at the end of the alignment. If *s* and *t* align in the opposite DNA complement, we instead link *s*_1_ to *t*_2_ and *s*_2_ to *t*_1_. We then perform within-contig clustering, which resolves variability in endpoint locations by clustering endpoints that refer to a similar part of the contig, meaning that they are (1) at most *d* bps from one another (*d* = 75 by default); and (2) refer to intervals in the same position with respect to the endpoint (to the endpoints’ “*right*” or “*left*”).

Next, operating across contigs, we merge clusters of endpoints that refer to the same side of pairwise alignments, using the links between endpoints that were described above. During this process we also resolve mosaic repeats, which, due to the intransitivity of pairwise alignments, may lead to incomplete identification of homology groups and erroneous copangraph structure (**Fig. S2**). Mosaic repeats are resolved by detecting endpoints that fall between larger alignments on one contig, translating that endpoint’s position to its corresponding position within the other contig of the alignment, and then splitting the larger alignment by introducing additional endpoints at the translated location (**Fig. S4**, **Supplementary Algorithm 3**). We iterate this process until all detected mosaic repeats are resolved (**Fig. S3)**.

We then use the clustered endpoints to define disjoint intervals that completely cover each contig without overlaps. This is done by computing, for every contig, consensus points from the clustered endpoints, each defining a shared boundary of a pair of contiguous intervals - one to the left of the consensus point, and one to the right - which are disjoint (**Supplementary Algorithm 3**). We then organize these disjoint intervals into homology groups based on the endpoints each consensus point summarizes. If a disjoint interval is involved in a set of pairwise alignments, then the consensus point defining its left bound will summarize a set of clustered endpoints representing pairwise alignments to their *right* (i.e., those involving the interval itself). Similarly, the consensus endpoint defining its right bound will summarize a set of clustered endpoints representing alignments to their *left*. Since the endpoints on either side of a pairwise alignment are clustered with those representing the same side of homologous intervals across other contigs and in other samples, the cluster assignment of these endpoints determines homology group assignment - therefore, we assign all (*right, left*)-bounded disjoint intervals with the cluster pair to the same homology group. Note that, disjoint intervals that are not (*right, left*)-bounded have no detected homology to any other intervals, and we therefore assign each to a singleton homology group (**Supplementary Algorithm 3**).

#### Tunable graph merging: graph construction

Copangraph construction (**Supplementary Algorithm 4:** `constructCopangraph`) starts by collapsing disjoint intervals of the same homology group into copangraph nodes. During node construction, one complement is arbitrarily defined as ‘+’ and the other ‘−’, and all sequences in the homology group are assigned a complement according to those definitions; if only one sequence is present in a node, its complement is defined as ‘+’ by convention. After we construct all nodes, we turn to constructing edges. We induce a copangraph edge between nodes *u* and *v* if: (1) there is an interval in *u* and an interval in *v* that are contiguous in some contig (in other words, whenever a contig is “split” between two nodes we induce an edge between them); (2) if there is an edge between an interval in *u* and an interval in *v* in any of the sample-specific sequence graphs (in other words, copangraph “inherits” edges from the single-sample graphs).

Edges in a copangraph are bi-directed, and reflect the relative orientation of the intervals in the nodes they connect. To briefly define bi-directed edges, we first define *a*^+^ and *a*^−^ as the forward and reverse complement of contig *a*. Then, the bi-directed edge 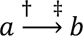 means that the concatenation *a*^†^*b*^‡^ exists in some genome. Because the sequence complement defined as ‘+‘ in each sample-specific sequence graph could be different than the complement defined as ‘+‘ in copangraph, when “inheriting” edges from single-sample graphs, their bi-directedness needs to be redefined such that it consistently refers to the same complement. For instance, let *a*_1_*, b*_1_ be contigs in sample 1 connected with the bi-directed edge 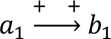, and *b*_2_*, c*_2_ be contigs in sample 2 connected with 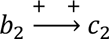, where *b*_1_ is in the opposite complement to *b*_2_. A copangraph with these contigs will have three nodes: 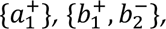, and 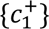. Note that because *b*_1_ and *b*_2_ are in opposite complement, *b*_2_ is in the reverse complement to that of its copangraph node. Therefore, the bi-directed edges need to be redefined before they are inherited into copangraph. We resolve this by multiplying the complement signs associated with sequences in copangraph nodes, which are synchronized across the dataset, with the sample-specific bi-directed edges. In this case, multiplying the signs in 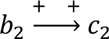 by the signs of 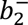 and 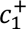 translates the edge into 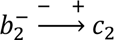, with a final graph of 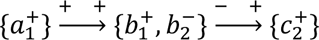.

Copangraph maintains the original sequence of all the intervals collapsed into a node, as well as the sample of origin for each contig and edge. We output copangraph in Graphical Fragment Assembly format^96^, with corresponding node-by-sample and edge-by-sample tables describing sample occurrence.

### Gut metagenomic sequencing and VRE culture results from liver transplant recipients

We performed metagenomic sequencing of 77 gut microbiome samples collected from a previously described longitudinal cohort of liver transplant recipients^14,50^. The study was approved by the IRB of Columbia University, approval number AAAM7704, and all participants provided informed consent. We used all samples which had a concurrent selective culture using chromogenic agar that indicated colonization by VRE, and which had an additional VRE culture result within 3-28 days^14,50^. The characteristics of the selected subcohort are detailed in **Table S1**. For the metagenomic sequencing, initial sample pre-processing, including stool homogenization and aliquoting, was performed using the epMotion automated liquid handling system (Eppendorf, Hamburg, Germany). Microbial DNA was extracted using the DNeasy 96 PowerSoil Pro QIAcube HT kit (Qiagen, Hilden, Germany, cat. 47021) following the manufacturer’s protocol. Extraction was automated using the QIAcube HT platform. Both a positive and negative control were included in each extraction batch. The ZymoBIOMICS Microbial Community Standard (Zymo Research, Irvine, CA, USA, cat. D6300) was used as a positive control, and biological grade water (Citiva, USA, cat. SH30538) was used as a negative control to monitor for potential contamination. Shotgun metagenomic libraries were prepared from extracted DNA using the Illumina DNA Prep kit (Illumina, San Diego, CA, USA, cat. 20060059) according to the manufacturer’s instructions. Libraries were sequenced using an Illumina NovaSeq 6000 with a loading concentration of 2 pM and a 2% PhiX spike-in control to a mean±std depth of 24.6±12.1 million reads.

### Metagenomic data processing

Metagenomic sequencing reads from each sample were quality filtered using Trimmomatic^97^ v0.39 in order to remove reads containing Illumina adapter sequences, leading or trailing bases with quality scores below 25, and reads shorter than 50 bases. Metagenomic reads were then aligned to the human genome (CHM13V2) and PhiX genome via Bowtie2 (ref. ^93^) v2.5.3. Any reads where either end mapped were subsequently removed. Microbial abundances were estimated using mOTUs3 (ref. ^61^) v3.1.0 and gene (functional) abundances using HUMAnN3 (ref. ^62^) v3.9.

### Copangraph visualization

For **Fig. 1**, we constructed a multi-sample copangraph from 50 of our gut metagenomic samples from these liver transplant recipients. In parallel, all MEGAHIT assemblies used for copangraph construction were binned with METABAT2 (ref. ^98^) v2.17 run in default settings, and each bin taxonomically classified with GTDB-TK^99^ v2.2.1. We visualized copangraph in Gephi^100^ v0.10.1, applying the ForceAtlas2 layout^101^. To visually represent sequence length in Gephi, we replaced each copangraph node with a chain of ceil(max_seq / 2500) nodes with an in-degree and out-degree of one, where max_seq is the longest sequence assigned to the node.

### Coverage metric

Coverage quantifies how accurately a set of contigs represents a ground-truth reference metagenome at the bp level. We first map all contigs to the ground-truth metagenome with Mappy v.2.27, which is a python wrapper of minimap2 (ref. ^95^). To avoid mapping artifacts, we: (1) only consider mappings if the alignment length is within contig length ± 0.02 * contig length; and (2) require the number of bp mismatches to be less than 0.02 * contig length. To quantify coverage for sequence-graphs, we use the sequences represented by graph nodes as the contig set. Note that the following accounts for the fact that assembly graph nodes represent one sequence, while copangraph nodes represent a set of sequences. Each bp of the reference metagenome is classified as a true positive if all the sequences mapping to it come from the same node. On the other hand, if no sequence maps to it, it is classified as a false negative. If the sequences mapping to it come from n > 1 nodes, we count one true positive and (n-1) false positives. Finally, if any sequence from the contig set fails to map to any genome in the reference, each of its bps contributes a false positive. For clarity, this would include a sequence that is in the same node with sequences that do map to the reference (i.e., the same node could contribute both true positives and false positives).

### Connectivity metric

Connectivity quantifies how accurately a sequence-graph represents variant information, either via assembly or graph connectivity, and as such is more suitable for sequence graphs. Like coverage, it uses a ground-truth reference metagenome. We identify variable positions using contig edges, as assemblers tend to end contigs in variable positions^9,32^. In some cases, however, such breakpoints will be generated due to low coverage, so we exclude breakpoints that do not have contigs mapped on both sides. Because MEGAHIT contigs contain overlaps of up to k bps (max k=141), we also cluster breakpoints that are a distance of ≤200bp and represent each cluster by the median position. We classify a breakpoint as a true positive if (1) a single contig completely maps across it; or (2) if two sequences from the graph map to the sides of the breakpoint and the nodes to which they are assigned are connected by an edge. Otherwise, it is classified as a false negative. To account for assembly artifacts, rather than requiring a pair of sequences to exactly flank a breakpoint, we instead require them to be in close proximity to the breakpoint, which we define as a maximum distance of 175bp (or k+34 for a default k=141). Finally, we check whether each edge is a false positive by pairing all the sequences represented by the two nodes it connects and checking whether they map to any genome in close proximity (defined as a distance of ≤175 bps between their closest start and end mapping coordinates). If no pairs map, the edge is classified as a false positive.

Connectivity has a number of intuitive outcomes for extremes in the assembly quality. For example, a tool able to construct a “perfect assembly” from input contigs, i.e., construct only complete genomes, scores a connectivity F-score of 1 because every breakpoint is a true positive and there are no false positives. On the other hand, a contig set, i.e., a sequence graph with no edges and where every node is a contig, has a recall of 0 because all breakpoints will be false negatives, but a precision of 1 since there are no edges. Furthermore, a graph can have an arbitrary number of edges and not be penalized, so long as the sequences it connects are contiguous. For example, a raw de Bruijn graph (where each node is a k-mer) would not be penalized for having an edge between overlapping k-mers.

### Assembly, coassembly with alternative methods for benchmarking

Across different benchmarks (**Figs. 3b,d-h**), we compared copangraph with MEGAHIT contigs, MEGAHIT assembly graphs, and MetaCarvel scaffold graphs. For MEGAHIT contigs, we used the final.contigs.fa produced by MEGAHIT^54^. For MEGAHIT assembly graphs, we used the graphs produced by megahit_toolkit^54^ v1.2.9 from the contigs output by MEGAHIT. We followed the recommended guidelines (https://github.com/voutcn/megahit/wiki), and constructed graphs using the contigs in the ./intermediate_contigs directory, choosing to use the largest k available. For coassembly, we pooled reads from all samples together (i.e., concatenated the .fastq files), and then coassembled contigs with MEGAHIT and constructed coassembly graphs with megahit_toolkit as described above. To construct single-sample MetaCarvel scaffold graphs, we followed the construction guidelines (https://github.com/marbl/MetaCarvel/wiki), and mapped paired-end reads back to MEGAHIT- assembled contigs using Bowtie2 (ref. ^93^) v2.5.3 in single-end mode, which we then merged and sorted by read identifier with samtools^102^ v1.18 into a final .bam file that we input into MetaCarvel^35^ version - d4b5408 alongside the contigs. For multi-sample scaffold graphs, we pooled reads by concatenation and proceeded similarly with mapping and MetaCarvel construction with the coassembled contigs.

### Simulated short-read coverage and connectivity benchmarks

For the simulated benchmark (**Fig. 3b,d**), we simulated short-read metagenomic data with CAMISIM^52^ v1.3-fcc198. We randomly selected ten short-read gut metagenomic samples each from our liver transplant cohort, each from a different patient. We computed a taxonomic abundance profile for each using MetaPhlAn4 (ref. ^81^) v4.1.0, which we provided to CAMISIM as a BIOM file. We ran CAMISIM in taxonomic profile-based community design mode using the default Hiseq2500 error profile of art_illumina^103^ v2.5.8 to construct simulated reference metagenomes, and simulated corresponding metagenomic samples to our ten selected samples with 80 million read-pairs each. To construct samples at different depths, we randomly sub-sampled reads-pairs from each of the samples at the depths of 1, 2, 5, 10, 20, 40, and 60 million read-pairs.

### Paired long-read coverage and connectivity benchmarks

For the paired short-read and long-read analysis (**Fig. 3e-h**), we used publicly available metagenomic sequencing data^56^, accessible under NCBI BioProject PRJNA754443, which includes 12 stool samples from 6 participants that were sequenced with both Illumina NovaSeq 6000 and Pacbio Sequel 2. Of the 12 stool samples, long-read data was available for 11 of them, and two failed to assemble by MEGAHIT within five days, leaving nine samples with which the benchmark analyses were undertaken. Short reads were processed as described above.

To construct long-read reference metagenomes for this benchmark we constructed long-read metagenome-assembled genomes (MAGs). Each long-read sample was assembled with MetaFlye^104^ v2.9.5-b1801, and the resulting assemblies were filtered to include only contigs longer than one million bps. To remove potential mis-assemblies, we used CheckM^105^ v1.2.3 to identify contigs with contamination > 5%. For each benchmark, we pooled the contigs across the samples it included (3-9 in **Fig. 3e,f**, one in **Fig. 3g,h**), clustered them a 99% ANI with dRep^43^ v3.5.0, and selected the contig representing with the highest completeness in each cluster as a representative MAG.

For the multi-sample benchmarks (**Fig. 3e,f**), the coassemblies of three and six samples were chosen at random. Multi-sample copangraphs, multi-sample MetaCarvel scaffold graphs, MEGAHIT coassembled contigs and coassembly graphs were constructed using the short-read data of each sample subset. For all benchmarks, coverage and connectivity were calculated with two differences: (1) Because it is unlikely that long reads represent the complete taxonomic content of the samples, the basepairs of unmapped sequences were not counted as false positives for coverage, and edges involving a node representing unmapped sequences were not counted as a false positives for connectivity; and (2) to distinguish between closely-related representative MAGs the number of bp mismatches allowed between a mapped sequence and the reference metagenome was changed to <0.001 * sequence length.

### Analysis of copangraph structure and homology threshold

For the analysis in **Fig. 4a**, we simulated ten populations of 25 genomes using Bacmeta^57^ v1.1.0, with each population initialized to a different seed genome with a length of four million bps. We evolved each population for 25,000 generations with an single-nucleotide variant mutation rate of 2✕10^-6^ and indel mutation rate of 2✕10^-7^. At every 1,000th generation we computed the average pairwise ANI between all 25 genomes in each population with FastANI^106^ v1.1.0. We then constructed copangraphs from each population at different homology threshold values, and quantified the number of nodes in each copangraph.

### Copangraph richness analysis

For the analysis in **Fig. 4b**, we assembled each of the ten simulated human gut metagenomes sampled at 20 million read-pairs with MEGAHIT, and constructed copangraphs for a range of sequence identity thresholds between 70-100%. For each assembly, we derived a contig-by-genome occurrence matrix by mapping the contigs of a sample against its corresponding reference metagenome using BLAST^107^ v2.15.0+. Partial and low quality mappings were excluded by only including mappings where the sequence identity was ≥98% and the alignment length was >98% and <102% percent of the contig length. Combining this occurrence matrix with the contigs whose intervals were represented in each copangraph node and the taxonomic profile provided by CAMISIM provided us a full taxonomic composition of each node. The species, genera, and family richness of each node was calculated by counting the number of unique taxa present in each node.

### Analysis of copangraph and coassembly graph properties

This analysis, shown in **Fig. 4c-f**, used the nine paired short- and long-read sequenced samples^56^. To obtain MEGAHIT coassemblies with different k-mer sizes, we constructed a coassembly over these samples using MEGAHIT with default settings, and then ran ’megahit_toolkit <*k>* ./intermediate_contigs/k<*k>*.contigs.fa > k<*k>*.fastg’ over each specified k-mer value used by MEGAHIT. Coverage was calculated as in **Fig. 3e**. To calculate the reference proportion in multi-genome nodes (**Fig. 4e,f**), we first map each sequence in the graph to the representative MAGs from all 9 samples, including only alignments where (1) alignment length is within sequence_length ± 0.02 * sequence length; and (2) the number of bp mismatches < 0.001 * sequence length. We use these alignments to define “multi-genome nodes” as nodes whose assigned sequences collectively map to more than one reference MAGs. We then sum the total number of bps in the reference sequence that are aligned to sequences in multi-genome nodes, avoiding double-counting bps from overlapping intervals. Dividing this count by the total length of the reference metagenome - sum of the length of each representative MAG - provides us the reference proportion in multi-genome nodes.

### Analysis of strain-level variation

We used the BacMeta^57^ v1.1.0 simulated genomes from **Fig. 4a** that had an average nucleotide identity of 97%. We then simulated sequencing data to 10x depth using the art_illumina^103^ v2.5.8 sequence simulator with the HiSeq error profile. This process was done in triplicate, using genomes produced using a different seed each time. Each population was then coassembled with MEGAHIT, and a coassembly graph was constructed for each intermediate_contigs/k<*k*>.contigs.fa with megahit_toolkit. Copangraphs were constructed by considering the simulated data from each genome as a separate sample. Copangraphs were constructed with sequence identity threshold ranging from 90% to 99.9%. For each graph, coverage F-score and the proportion of reference in multi-genome nodes were calculated using the reference genomes.

### Resource benchmarks

For all tools, we calculated runtime and peak memory usage as the walk clock time and maximum resident set size output from the `/usr/bin/time -v` UNIX executable. MEGAHIT and metaSPAdes coassemblies were run with 64 threads and default settings. For copangraph, the first stage of hybrid coassembly processes each sample separately, and we used 8 threads on a similar machine. To allow comparison with MEGAHIT and metaSPAdes^55^ v.4.2.0, we emulated parallelism on a 64 thread machine with the following equation, 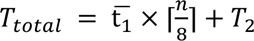 where 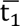 is the average first stage runtime for processing each sample with 8 threads, and *n* is the total number of samples. 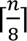 assumes 8 samples can run in parallel - which caps the thread usage to 64 - and ensures an additional 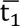 time when *n*% 8 ≠ 0. *T*_2_ is the runtime for the second stage of hybrid coassembly. The peak memory usage for copangraph was calculated as the peak memory usage reached by the second stage or any single sample run of the first stage.

### Analysis of gut metagenomic samples from a study of colorectal cancer

We analyzed publicly available metagenomic data from Yu et al.^59^, selected as the largest study in a previous meta-analysis^64^, which we downloaded from the European Nucleotide Archive accession number PRJEB10878. Yu et al. collected stool samples from 128 individuals. The study included 75 patients with colorectal cancer (15 stage I, 21 stage II, 34 stage III and 4 stage IV) with a median age of 67 years, of which 26 were female; as well as 53 participants designated as controls with a median age of 62 years, of which 21 were female. We performed QC, adapter trimming, and Human and PhiX filtering as detailed above. One post-processed sample was removed due to low sequencing depth (<0.5M read pairs) resulting in 127 samples with a mean±sd of 2.7×10^7^±4.9×10^6^ read pairs.

### Analysis of vaginal metagenomic samples from a study of colorectal cancer

We analyzed publicly available metagenomic data from the MultiOmic Microbiome Study: Pregnancy Initiative (MOMS-PI) PTB case-control study^60^, which we obtained from dbGaP (study no. 20280; accession ID phs001523.v1.p1). We performed QC, adapter trimming, and Human and PhiX filtering as detailed above., and selected samples with over 0.5M read pairs collected between 16 and 24 weeks of gestation. In case of multiple samples available per patient, we selected the latest one. This resulted in 100 samples with a mean±sd of 6.4×10^6^±1.2×10^7^ read pairs. 29 of these samples were from individuals who eventually had spontaneous preterm birth, and 71 from those who eventually delivered at term.

### Calculation of distance metrics between samples in copangraph

To select abundant-yet-variable parts of the pan-metagenome, we first constructed a samples-by-nodes abundance matrix from copangraph by calculating the reads per kilobase million (RPKM) of each contig in each sample, and then populating each entry of the matrix with 0 if the corresponding sample does not have a contig interval occur in the corresponding node, or to the RPKM value of the contig if it does. We then filtered this matrix, retaining all nodes with between 10-40% of the entries with RPKM≥50. Euclidean distances were compute directly on this filtered matrix, and the Jaccard distance was calculated on the filtered matrix after binarization.

### VRE colonization trajectory and prediction benchmarks

For the VRE colonization analysis (**Figs. 7,S1**), we defined persistence and clearance labels using the closest culture result to the metagenomic sample collection date within the subsequent 3-28 days window^14^. For the prediction model based on clinical covariates, we used age, sex, BMI (kgm^-2^), MELD score, Child-Pugh score, number of days the sample was taken post transplant, total number of antibiotic course days up to the sampling date, and the following etiologies of liver disease for each patient each encoded as a binary “yes/no” column: hepatitis C infection, hepatitis B infection, alcohol-related liver disease, non-alcoholic fatty liver disease, hepatocellular carcinoma, and an “other” variable capturing various additional etiologies such as polycystic liver disease or genetic/metabolic diseases. To construct the copangraph feature table, we concatenated the sample-by-node and sample-by-edge occurrence tables output by copangraph together to produce a single feature table, and then removed features that were identical to one another (retaining a randomly selected one).

For each feature type, we trained LightGBM^108^ v.4.1.0 classifiers in a nested cross-validation setting, using 10 outer folds to evaluate performance and 5 inner folds to optimize hyperparameters, while always keeping samples from the same patient in the same fold. We removed features with less than 20% non-zero entries, and performed additional feature selection within the nested cross-validation framework by training a lasso^109^ model and retaining only those with non-zero coefficients. We searched over an identical hyperparameter space (**Table S2**) for copangraph, mOTUs3, spacegraphcats and HUMAnN3 classifiers, allowing exploration of up to 1000 iterations of randomly selected hyperparameters. The model based on clinical covariates was trained with the following changes: (1) missing values were imputed within the nested cross-validation setting using a k-nearest neighbor model (k=5); (2) because of the number of features, we did not use feature selection. We concatenated the test predictions from each outer fold to compute the ROC curve for feature type.

For HUMAnN3, we evaluated both the gene and gene-by-organism abundance tables, and reported results for the former, which outperformed the latter. For spacegraphcats, we ran the first and second stage from the Reiter et al.^53^ pipeline on all 77 metagenomic samples. In the first stage, we set min_sample_frac to 0.8, within the recommended range, after the default value of 1 yielded no shared genomes. We ran the second stage with the default setup, which yielded 33 <*genome*>_abund_pruned.tsv files, each describing “graph piece abundances”^53^ for one genome. We then concatenated these together to construct a sample-by-“graph piece abundance” feature table which was used for prediction. Spacegraphcats was run only for the liver transplant dataset due to prohibitively long run times.

### Copangraph-based comparative metagenomics

We randomly selected one of the copangraph-based classifiers from the outer cross-validation folds, and evaluated its feature importance using SHAP^110^ v0.45.0. For each of the top five features (by absolute SHAP value), which is either a node or an edge, we extracted the surrounding copangraph subgraph up to a distance of 10 nodes from the feature. If a node had a degree > 20, we include it but not its immediate neighbours, unless they were already added to the subgraph.

We annotated contigs from all samples with Prokka^111^ v1.14.6, and used them to annotate the subgraphs shown in **Fig. 7**. For the prophage-containing subgraph (**Fig. 7g**), Prokka annotated only hypothetical proteins. We therefore annotated the contigs present in this subgraph also with geNOMAD^72^ v1.11.0 which provided annotations of phage-related proteins and prophage elements. We visualized each subgraph in Bandage^112^ v0.8.1, setting the node width to the number of samples in each node. In Bandage, to aid visualization, we reduced the subgraphs to a closer vicinity of the feature using the “around nodes” setting, and merged nodes in unambiguous chains. Except for the prophage graph (**Fig. 7g**), in which we wanted to demonstrate variable integration, nodes representing just one sample were removed.

### Experimental validation of VRE strain variability

To confirm copangraph “bubbles” represented strain variability, we sequenced 23 VRE strains from the same set of stool samples, each from a different patient, using chromogenic agar as published previously^14^. We performed nanopore long-read sequencing as described previously^113,114^. Briefly, we used the Rapid Barcoding Kit (SQK-RBK004 or RBK110.96) to prepare multiplexed DNA libraries for nanopore sequencing on R9.4.1 flow cells using the GridION (Oxford Nanopore Technologies).

Basecalling and demultiplexing were performed by MinKNOW (Oxford Nanopore Technologies); reads were then trimmed using Porechop^115^ v0.2.1. We assembled the resulting data using Flye^116^ v2.9.5. Sequences belonging to each branch of the “bubbles” were then mapped against the resulting assemblies with BLAST^107^ v2.16.0+. An isolate was considered to “belong” to a branch if its sequence mapped with >99.5% sequence identity. One of the 23 isolates did not map to either branch of any of the bubbles in **Fig. 7c,e**.

## Data availability

Individual-level clinical data, clearance and persistence labels, and sequencing data will be made available via the NCBI Sequencing Read Archive (SRA) (accession number PRJNA1297925) after filtering any human-derived sequences. Paired short- and long-read sequencing data^56^ is publicly available from NCBI, accession PRJNA754443. Metagenomic data and cohort metadata from Yu et al.^59^ is available from the European Nucleotide Archive (ENA), accession number PRJEB10878. Data from the MOMS-PI preterm birth case-control study^60^ is accessible from dbGaP (study no. 20280; accession ID phs001523.v1.p1).

## Code availability

Copangraph is available at https://github.com/korem-lab/copangraph. Code to generate all figures is available at https://github.com/korem-lab/MANU_copangraph.

## Acknowledgements

We thank members of the Korem and Uhlemann groups for useful discussion, as well as David Knowles and David Zeevi for discussions and advice. We are grateful to all investigators and participants involved in the generation of data used in this study. This study was supported by the Program for Mathematical Genomics at Columbia University (T.K), R01AI183668 (T.K. & A.-C.U.) and R01HD106017 (T.K.).

## Supplementary Figures

**Supplementary Figure 1.**
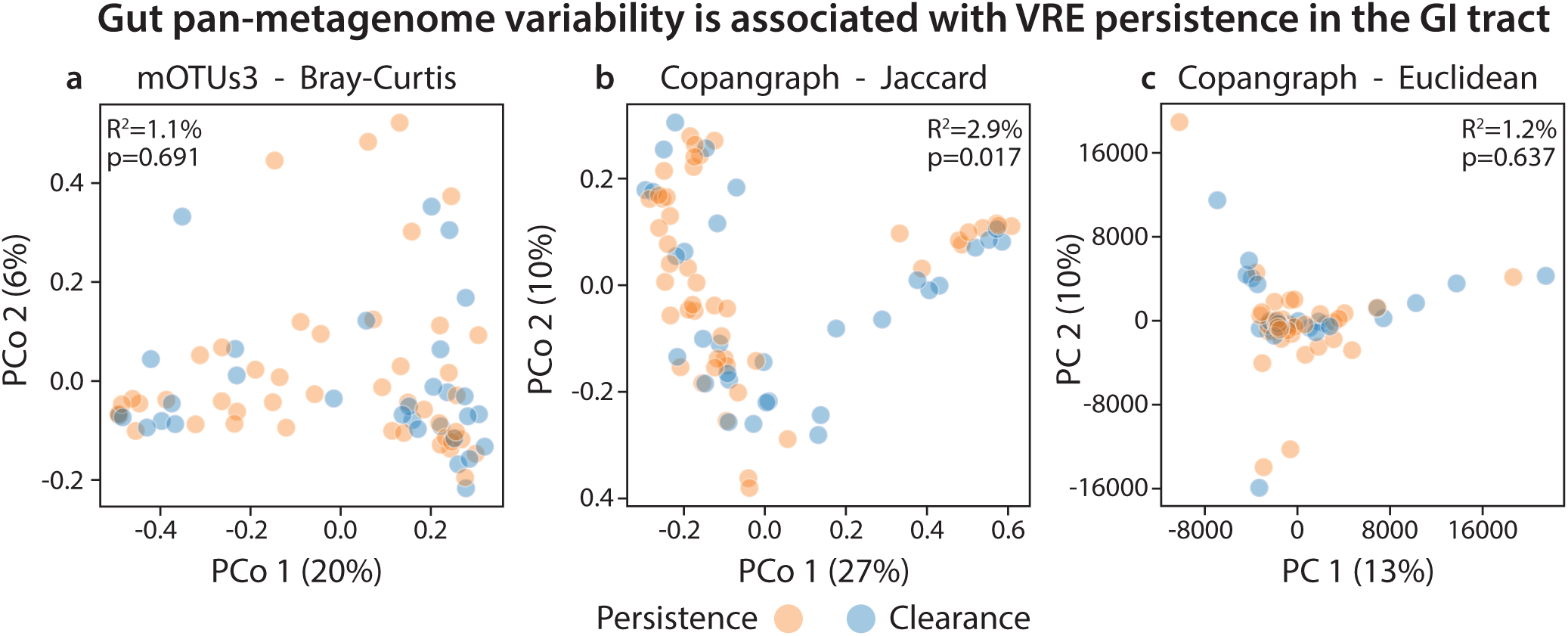
Copangraph represents metagenomic variability in a longitudinal cohort of liver transplant recipients. **a-c,** Shown are the first two PCos using Bray-Curtis dissimilarity calculated using species abundances (a), Jaccard distances calculated from copangraphs (b), or Euclidean distances calculated from copangraph (c). ADONIS^65^ explained variance and p value is displayed on each panel.

**Supplementary Figure 2.**
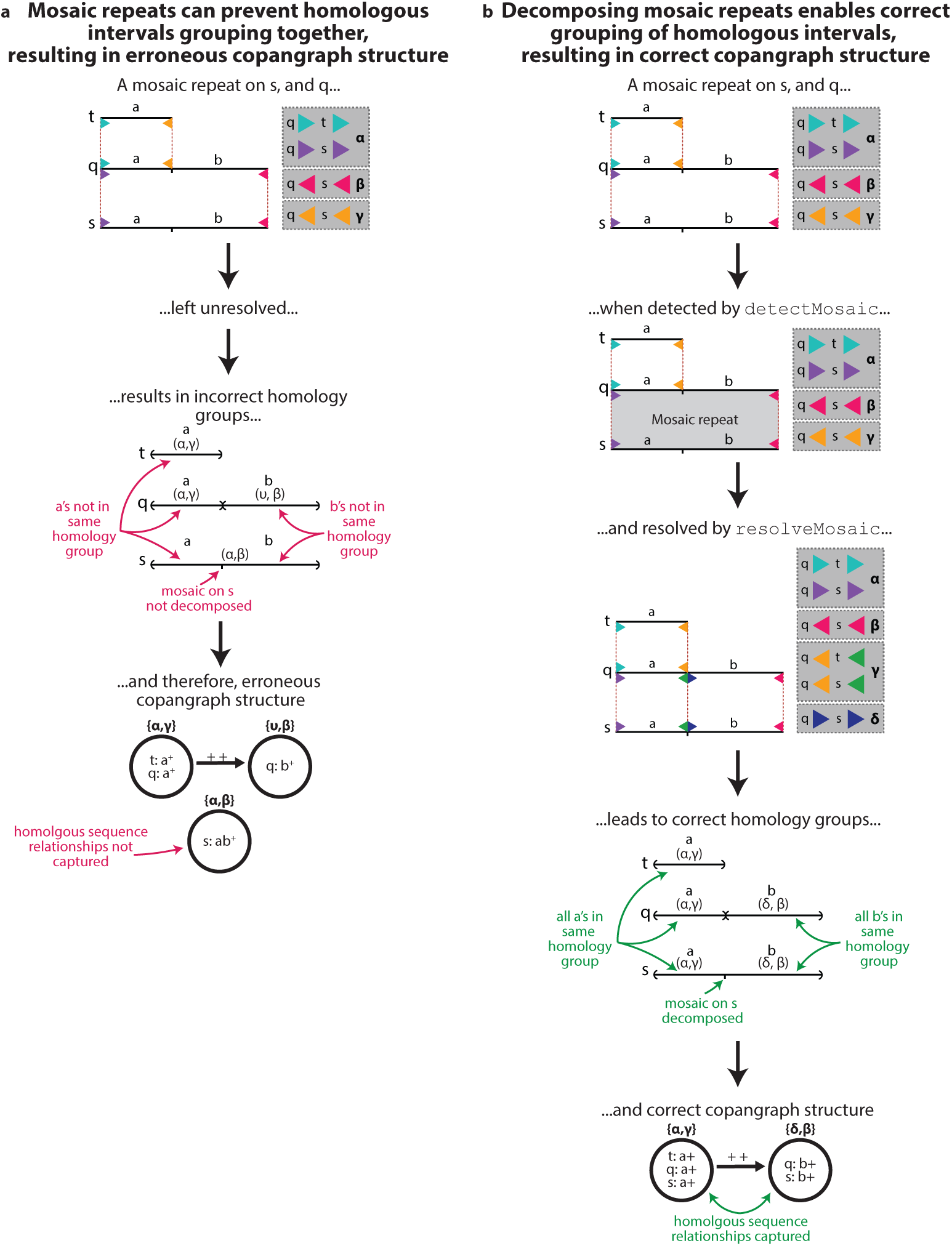
An example of the impact of an unresolved and resolved mosaic repeat on the structure of copangraph.

**Supplementary Figure 3.**
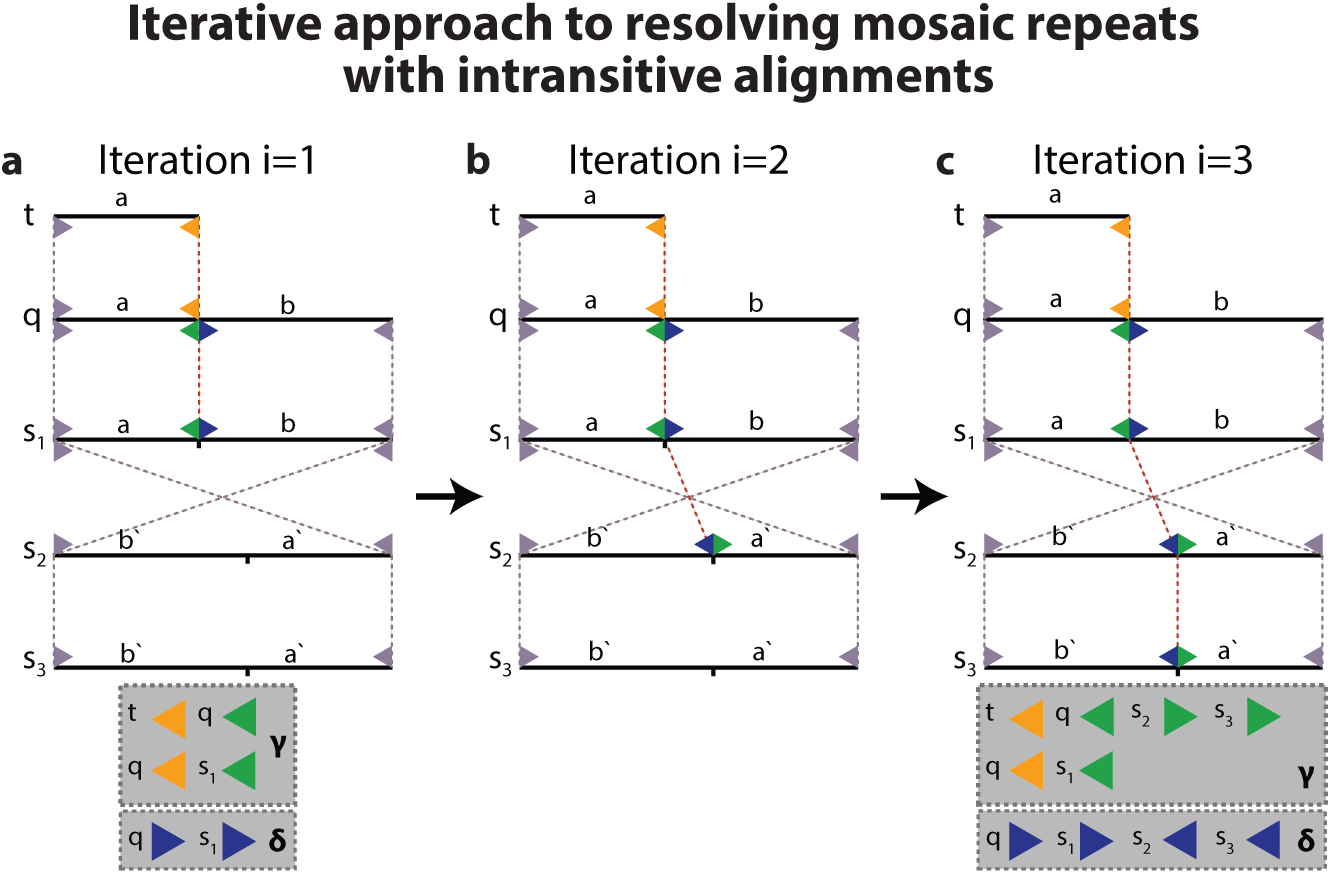
Iterative alignment splitting to convergence resolves mosaic repeats under intransitivity. **a,** A mosaic repeat 𝑎𝑏 occuring in 𝑞, 𝑠_1_, 𝑠_2_, 𝑠_3_ with 𝑎 repeating independently in 𝑡. 𝑎’ and *b*’ are the reverse complement of *a* and *b*. The alignments are intransitive - only three of the six possible pairwise alignments between *q, s*_1_*, s*_2_*, s*_3_ are present. One iteration of resolving mosaics introduces the green and blue endpoints on *q* and *s*_1_ between *a* and *b* in both contigs, thereby decomposing them into the individual repeating units. After this iteration, all endpoints to the right of *a* (yellow and green endpoints) are assigned to cluster **γ**, while those to the left of *b* (blue endpoints) are assigned to cluster **δ**; cluster assignments of the gray endpoints at the ends of the contig are not shown. **b**, Introducing the endpoints on *s*_1_ has revealed another mosaic repeat on *s*_2_which needs to be resolved by a second iteration. New points on *s*_2_ are introduced and correctly clustered, respecting the complement switch between *s*_1_ and *s*_2_. **c**, Another round of mosaic repeat resolution is needed for *s*_3_, after which no more mosaic repeats are detected.

**Supplementary Figure 4.**
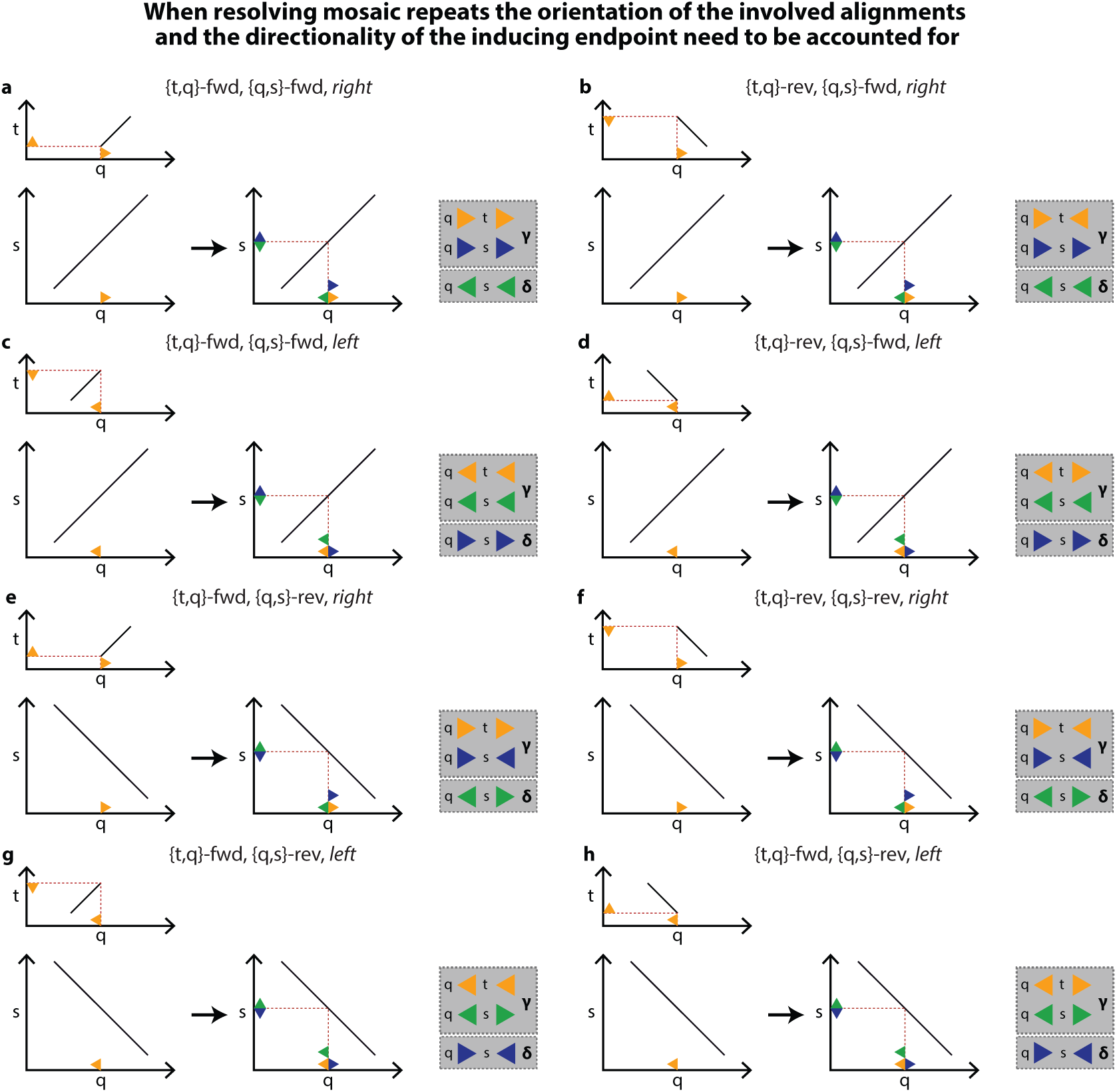
Correct mosaic repeats resolution depends on the direction of the inducing endpoint and the orientation of the mosaic alignment. A mosaic repeat is depicted in which an alignment between contigs *t* and *q* partly overlaps a larger alignment between contigs *s* and *q*, while no alignment is present between *s* and *t*. Alignments are shown as dotplots. An endpoint on *q* (yellow) at the bound of the {*q, t*} alignment is used to detect the mosaic repeat on *q* and introduce new endpoints (blue and green; added in plots to the right of the black arrow) that propagate the mosaic repeat into *s*. Each plot a-h shows a different combination of: (1) the sequence complement in which *t* and *q* align; (2) whether the interval common to *q*, *s*, and *t* is to the *left* or *right* of the initial endpoint; and (3) the sequence complement in which *q* and *s* align. In all cases, to correctly decompose the mosaic repeat, endpoints representing the two repeating intervals (one repeating on *q*, *s*, and *t*, the other repeating on just *q* and *s*) should be assigned to separate clusters (gray boxes).

## Supplementary Tables

**Supplementary Table 1.**
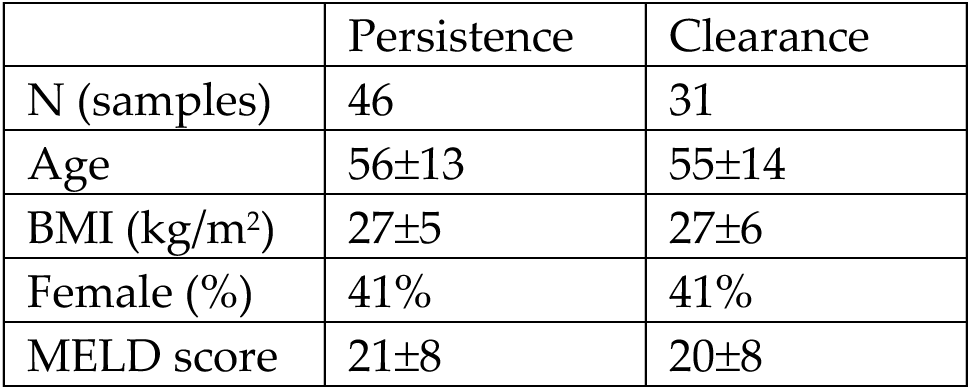
Liver transplant cohort characteristics. Shown is mean±std.

**Supplementary Table 2.**
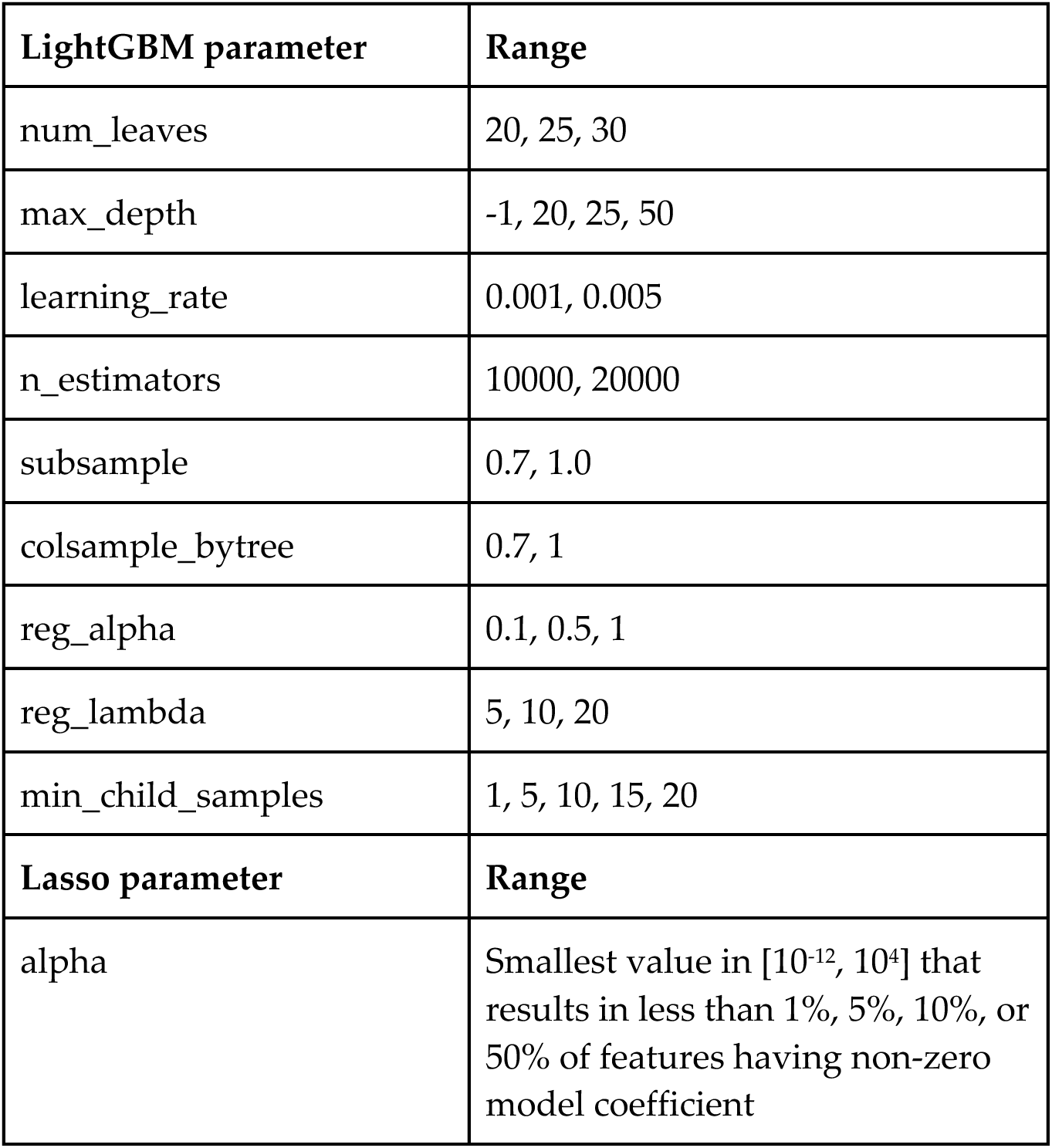
LightGBM hyperparameter space for classification, and lasso hyperparameter space used for feature selection. Feature selection was not used for the classifier based on clinical covariates.

## Supplementary Algorithm 1: buildSampleSequenceGraph

buildSampleSequenceGraph builds a single-sample sequence graph 𝒢(*V,* ℰ) from a set of paired-end sequencing reads ℛ. We begin by assembling a set of contigs 𝒞 from the reads using a separate assembler (line 1). We then align the reads to the contigs to construct **RdAlns**, a lookup table which, given an interval [*a* : *b*] on the contig *c* and a complement ( *forward* or *reverse*), returns all reads in ℛ that align to *c*[*a* : *b*] in the specified complement (line 2). Based on the mapping information in **RdAlns**, we calculate the mean and standard deviation of the paired-end fragment length *𝜇* and *𝜎*, as well as the average read length *𝜌* (line 3).

### Algorithm

buildSampleSequenceGraph(ℛ *, d*)

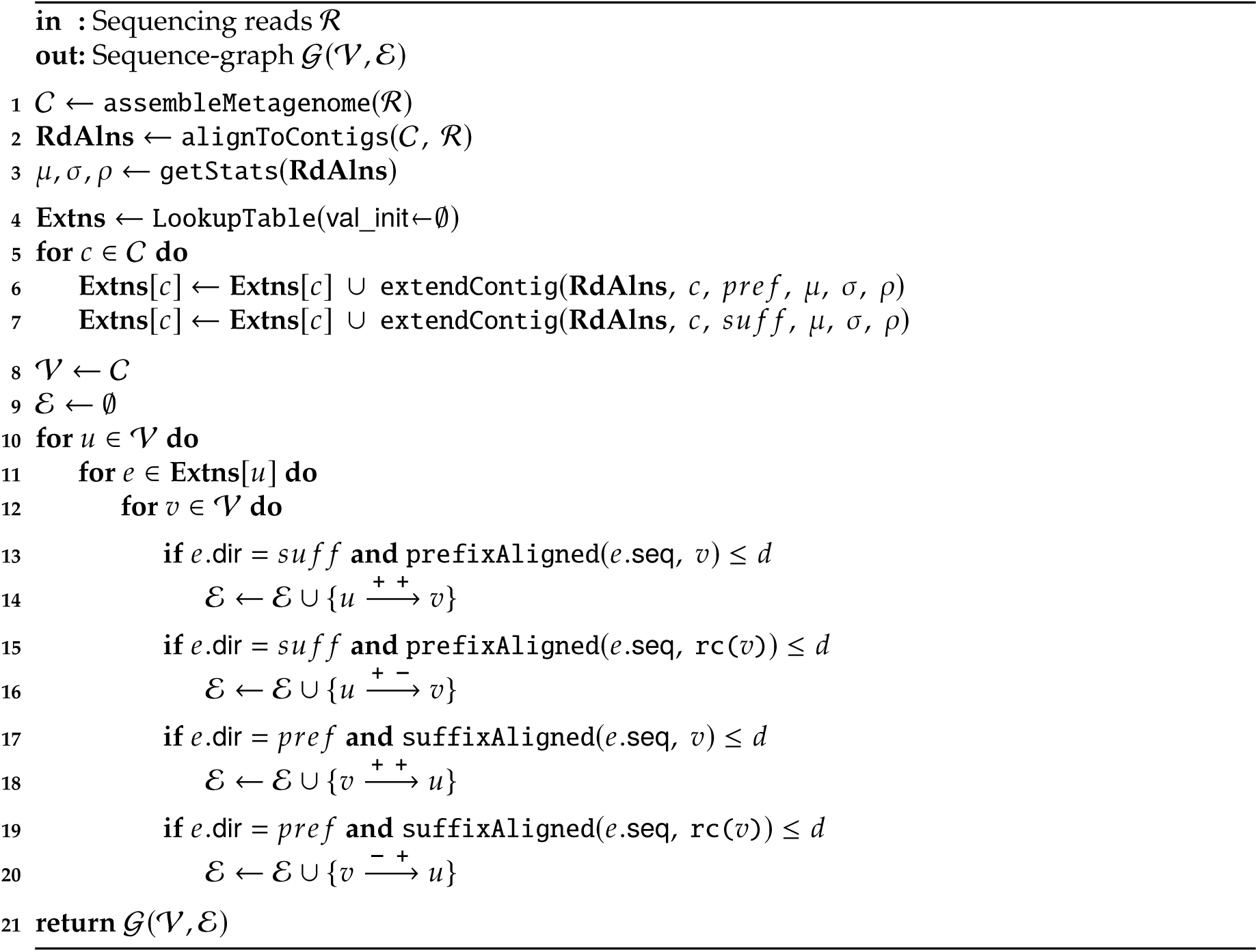

We then iterate over each contig *c* ∈ 𝒞 (line 5), and with extendContig (**Supplementary Algorithm 2**), assemble the genomic region immediately upstream or downstream to its prefix and suffix. We then add these “extensions” to a contig-specific extension set **Extns**[*c*] (lines 6-7). The nodes of the sequence-graph *V* are set to the contigs 𝒞 (line 8), and, for the remainder of this section, hold the same meaning.

We now use the extensions in **Extns** to construct the edges ℰ, by detecting overlaps between them and the other nodes (lines 10-20). Iterating over each node *u* ∈ *V* (line 10), we iterate over its set of extensions **Extns**[*u*] (line 11). Each extension *e* ∈ **Extns**[*u*] is a named tuple, with *e.*seq set to the sequence assembled by extendContig, and *e.*dir set to either *pre f* or *su f f*, depending on whether *e* was extended from the prefix or suffix of *u*.

We then iterate over each node *v* ∈ *V* and check whether the extension *e* aligns to the prefix or suffix of *v* with sequence identity greater than *d* (*d* = 99.5% by default), and with no alignment gaps (lines 13, 15, 17, and 19). Such a match indicates that *u* and *v* are contiguous in some genome, and should therefore be connected in 𝒢. We therefore construct a bi-directed edge between *u* and *v*, with an orientation that depends on whether: (1) *e*derives from the prefix or suffix of *u*; (2) *e* aligns to the prefix or suffix of *v*; and (3) *e* aligns to *v* in the forward or reverse complement (rc(*v*)) (lines 13-20). Letting *c*+ and *c*− be the forward and reverse complement of a contig *c*, a bi-directed edge 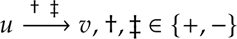, means that the sequence *u*†*v*‡ exists in some genome. We therefore construct a bi-directed edge between *u* and *v* specifying the complement of *u* and *v* that, when concatenated, correctly represents the genome and add the edge to ℰ (lines 13-20). Note that lines 13-20 ignore prefix-prefix and suffix-suffix matches (four of the eight possible cases defined by the logic in lines 13-20) as these do not describe valid connections between contigs.

## Supplementary Algorithm 2: extendContig

Assemblers typically terminate contigs at branch points, after which multiple sequences may follow. The prefix-suffix matches used by buildSampleSequenceGraph to connect nodes rely on extendContig, which uses paired-end information from the same sample to assemble the genomic region(s) immediately upstream or downstream to a contig’s prefix or suffix, which we term an “extension”. extendContig performs a recursive depth-first assembly, assembling each of the multiple sequences that may follow a contig. Each iteration of the algorithm considers the addition of one character to an extension. All possible extensions are considered if more than one DNA base is possible.

When determining possible bases for extension, extendContig considers only reads that are very likely to originate from the region currently being assembled. This is achieved using paired-end information: we consider read pairs in which one end already maps to the contig being extended (“mapped read end”), and consider the other end for extension (“extending read end”). To collect the set of mapped read ends, *mapped* (lines 1-10), we consider the contig *c* undergoing extension, as well as *𝜇* and *𝜎*, the average and standard deviation of the paired-end fragment length. We calculate a window on *c* that is centered *𝜇* characters away from the current extension edge, at the start or end of *c*, in which relevant mapped read ends are expected to map. To further enrich for reads over the extension edge, we also select reads mapped in the relevant complement (lines 4, 9). We then extract the *query*, a *k* = 31-mer at the current extension edge (lines 5, 10).

### Algorithm

extendContig(RdAlns, *c*, *dir*, 𝜇, 𝜎, 𝜌, *q*, *i* = 0, 𝒳 = ∅)

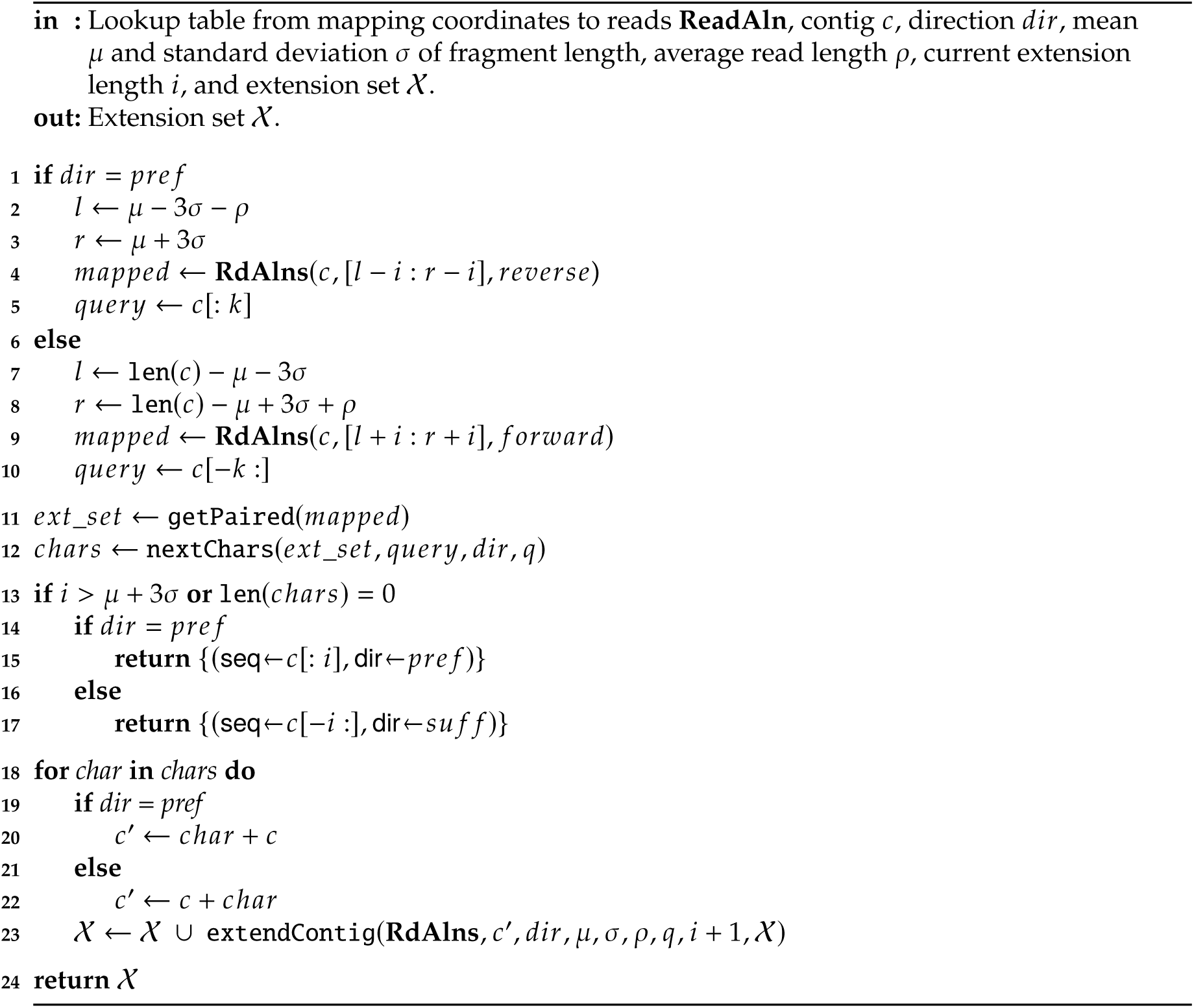

Next, we gather the extending read ends, *ext*_*set*, that are pairs of the reads in *mapped* (line 11). Using *query*, nextChars (line 12) then locally aligns the reads in *ext*_*set* to one another, and examines the aligned bases immediately following *query* in the direction of extension, considering only those with phred-quality ≥ *q* (default *q* = 30). The set of bases passing the filter is returned to *cℎars* (line 12). *cℎars* represents the next possible bases to append to the extension (between zero and four).

We terminate extension when there are no additional bases in *cℎars* or if the length of the extension, measured by *i*, is greater than the average fragment length plus three standard deviations, at which point little paired-end information remains to inform the extension process (line 13). Upon termination, the sequence of the extension is returned in a named tuple, along with the direction of extension (lines 14-17).

If extension did not terminate, we iterate over each base in *cℎars* (line 18), concatenate *cℎar* to the contig to produce a new contig *c*^′^ (lines 19-22) and then recursively call extendContig on the new contig, extending in the same direction and incrementing *i* by 1 (line 23). 𝒳, the set storing the extensions, is updated after each call to extendContig returns (line 23) and is returned on line 24.

## Supplementary Algorithm 3: createHomologyGroups

Tunable graph merging, the process by which we merge the individual sequence graphs produced by constructAssemblyGraph into a multi-sample copangraph, begins with createHomologyGroups. The contig sets from all samples, 𝒞_1:_*_𝑆_*, are first pooled together (line 1) and a minimizer-based co-linear chaining algorithm is used to detect pairwise alignments among them, keeping only alignments with sequence identity above the homology threshold *𝜏*and length longer than *l* (line 2).

### Algorithm

createHomologyGroups(𝒞1:𝑆, 𝜏, *l*, *d*)

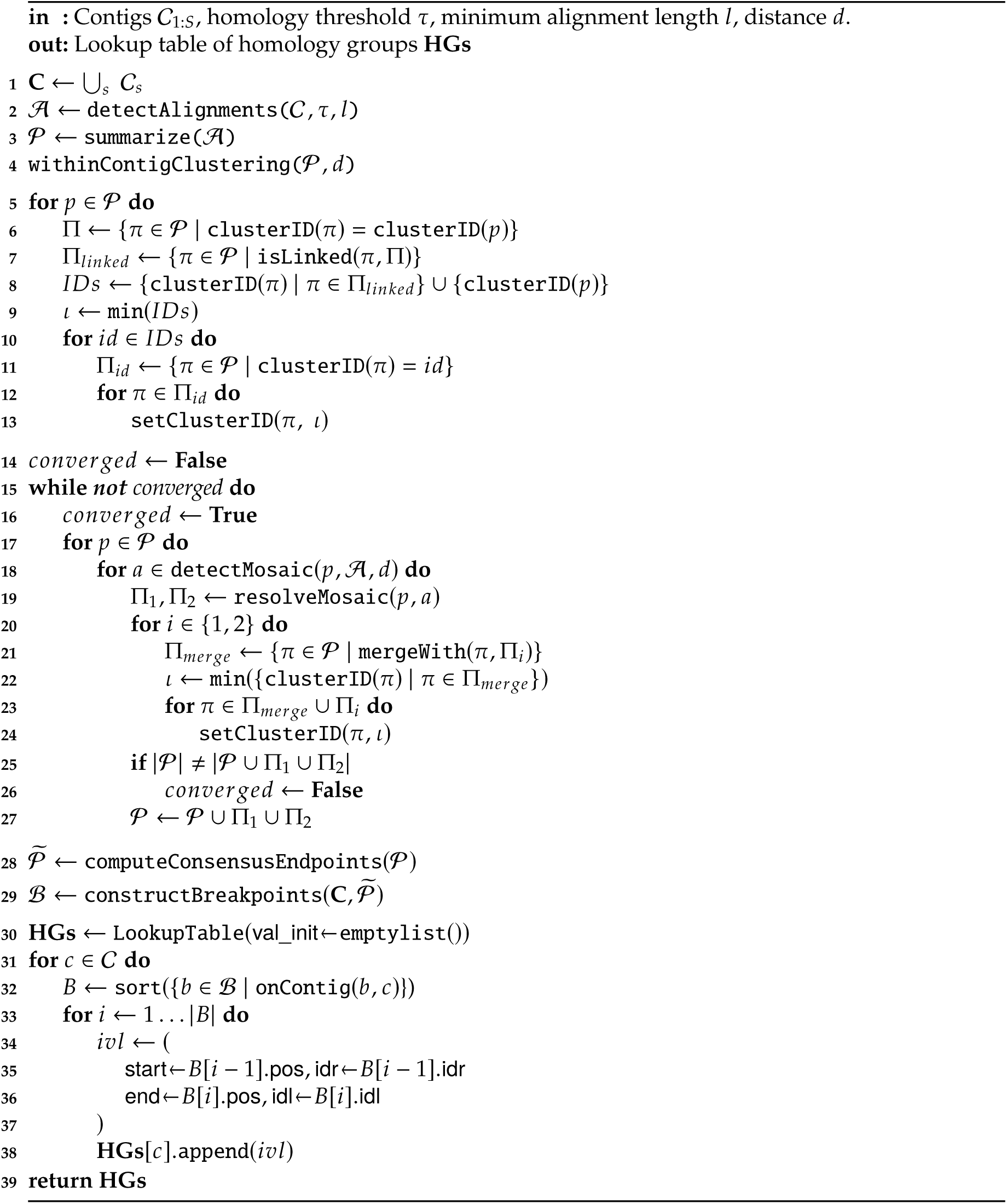

The alignments 𝒜 are summarized by their endpoints (line 3). Letting *s*[*i* : *j*] and *t*[*k* : *l*] be aligned intervals from two contigs, two endpoints are generated for *s*: *s*_1_ = (*i, rigℎt*) and *s*_2_ = (*j, le f t*), and two for t: *t*_1_ = (*k, rigℎt*) and *t*_2_ = (*l, le f t*). “*rigℎt*” and “*le f t*” specify whether the alignment is to the right or left of the endpoint (i.e., the alignment is to the *rigℎt* of *t*_1_ regardless of whether it is in the forward complement of *t*, that is, from *k* to *l*, or whether it is in the reverse complement, from *l* to *k*). The alignment itself is represented by links between the endpoints: if the intervals align in the forward complement, *s*_1_ links to *t*_1_, and *s*_2_ to *t*_2_; if the alignment is in the reverse complement, *s*_1_ links to *t*_2_ and *s*_2_ to *t*_1_.

The endpoints in 𝒫 now undergo withinContigClustering, a fast and greedy clustering algorithm that iterates over endpoints in order of their position along the contig, and merges consecutive endpoints if they have the same interval position (*rigℎt* or *le f t*) and if the distance between them is ≤ *d* (*d* = 75bps by default). All endpoints in the same cluster receive the same ID, retrievable with clusterID(*p*) (line 4).

Next, we perform between-contig clustering, by merging clusters of endpoints that refer to the same side of alignments between homologous intervals. As mentioned above, such endpoints are linked according to alignment complement. We define the function isLinked(*p,* Π) which returns true if endpoint *p* is linked to any endpoint in the set of endpoints Π ⊆ 𝒫, and false otherwise. Iterating over all endpoints *p* ∈ 𝒫 (line 5), we first construct Π, the set of all endpoints *p* is clustered with (line 6). These are initially all on the same contig, the result of within-contig clustering. We then construct Π*_linked_*, the set of all endpoints in 𝒫 that are linked to at least one endpoint in Π (line 7). The clusters to which all endpoints in Π*_linked_* and *p* are assigned are then merged into a single cluster, by assigning all of them the minimal cluster ID, *𝜄* (lines 8-13). By iterating through all the endpoints, we ensure single-linkage clustering of all alignment-linked within-contig clusters.

The goal of between-contig clustering is to eventually group together all homologous intervals across the different contigs. This, however, does not happen in some cases involving mosaic repeats, due to the intransitivity of the alignment process. A mosaic repeat is a concatenation of sequences, e.g., *ab*, that repeats in the data both together and as separate sequences. The mosaic repeat *q* = *ab* is “composed” of the individual repeating units *a* and *b*. An instructive example of a mosaic repeat involving intransitive alignments is seen in **Fig. S1**, in which contig *t* is composed of *a*, and contigs *q* and *s* are composed of *ab*. Ideally, following between-contig clustering, four clusters of endpoints will emerge: (1) connecting the endpoints at the start of *t*, *q*, and *s*; (2) connecting the end of *t* with endpoints between *a* and *b* in both *q* and *s*; (3) connecting the start of *b* in *q* and *s*; and (4) connecting the end of *q* and *s*. However, alignments are not transitive; and so, although *t* aligns to *q*, and *q* aligns to *s*, *t* and *s* do not align. Unless such mosaic repeats are resolved - decomposed into their individual repeating units - homologous sequences may not collapse into nodes, resulting in an “undercollapsed” copangraph (**Fig. S1a**). In the example, because *q* and *s* are not resolved into their *a* and *b* units, *s* is eventually assigned to a node separate from *t* and *q* despite their shared homology.

To address this issue, we: (1) detect mosaic repeats; (2) resolve them by introducing new endpoints in 𝒫 which “decompose” mosaics into their individual repeating units; and (3) correctly cluster the new endpoints to ensure the detection of homologous intervals and correct construction of homology groups. **Fig. S1b** provides an example of how these three steps resolve a mosaic repeat. Because the introduction of new endpoints in step (2) may lead to the detection of new mosaics, we iterate the process of mosaic repeat resolution until convergence, which is reached when no new endpoints are added to 𝒫 (lines 14-27). **Fig. S2** provides a visual example of the need to iterate mosaic repeat resolution to convergence.

To detect mosaic repeats, we define the function detectMosaic(*p,* 𝒜*, d*), which returns all alignments in 𝒜 which cover the endpoint *p* with one of the aligned intervals, so long as *p* is a distance ≥ *d* from the interval’s bounds (*d* = 75 by default). Considering the example in **Fig. S1b**, the {*q, s*} alignment contains a mosaic repeat, because the sub-interval *a* independently repeats in the data at *t*. In this case, the {*q, t*} alignment would generate an endpoint between *a* and *b* on *q*; detectMosaic will then return the {*q, s*} alignment, identifying it as covering a mosaic.

We iterate over all endpoints *p* ∈ 𝒫 (line 17), and use detectMosaic to check mosaics, i.e., alignments covering *p* (line 18). Letting, *a* = (*q*[*i* : *j*]*, s*[*k* : *l*]) be a mosaic alignment covering an endpoint *p* on contig *q*, we construct new endpoints on *q* and *s* by calling the function resolveMosaic(*p, a*).

In resolveMosaic, we first translate *p*’s position on *q*, *q*_*pos*, to its corresponding position on *s*, *s*_*pos*, using the aligned anchors - identical kmers shared between *q* and *s* ordered by their alignment positions - in *a* (line 1). We then construct two new endpoints on *s* with opposite directions to one another (line 2), and a new endpoint on *q* with opposite direction to *p* (lines 4 and 10). These new endpoints “propagate” the mosaic structure detected within *q*[*i,j*] into *s*[*k* : *l*], where it may be missing due to alignment intransitivity, and generate endpoints that decompose the structure into independently repeating units, namely *q*[*i* : *q*_*pos*]*, q*[*q*_*pos* : *j*], and *s*[*k* : *s*_*pos*]*, s*[*s*_*pos* : *l*].

We now need to integrate these new endpoints into the cluster structure over 𝒫 in order to construct the homology groups correctly, which is done by assigning them to the cluster representing the same side of the repeating region they refer to. We determine the correct assignment into two sets, Π_1_ and Π_2_, based on both the interval position of *p* and the complement (forward or reverse) of *a* (lines 3-14). **Fig. S3** provides a visualization of the correct organization and clustering of the new endpoints for each possible case in lines 3-14. Π_1_ and Π_2_ are then returned back to clusterHomologyGroups (line 15).

The endpoints returned from resolveMosaic in the Π*_i_*sets are already connected across contigs; so next, we need to check if they could be connected within their contigs. Therefore, for each Π*_i_* set (line 20), mergeWith identifies endpoints on the same contig, whose interval is in the same direction, and that are no further than *d* bps apart from an endpoint in the set (line 21). If such endpoints are found, we merge their respective clusters (22-24). We next check whether the new endpoints in Π_1_, Π_2_ increased the total number of unique endpoints in 𝒫 (lines 25-27). If they did, we continue with an additional round of mosaic detection (line 26). If after a complete iteration over 𝒫 no new endpoints are added, the while-loop exits. Finally, we summarize each set of clustered endpoints from the same contig to a consensus endpoint. However, because different regions from the same contig might end up in the same cluster, we once again cluster these endpoints according to orientation and position, as in withinContigClustering. We then compute a consensus endpoint, *p̃* = (median({*p*_0_*, …, p_n_* }), dir(*p*_0_)) over each set of clustered points *p*_0_, · · · *, p_n_* . This yields 𝒫^∼^, the set of consensus endpoints, which retain their clusterID.

### Algorithm

resolveMosaic(*p*, *a*)

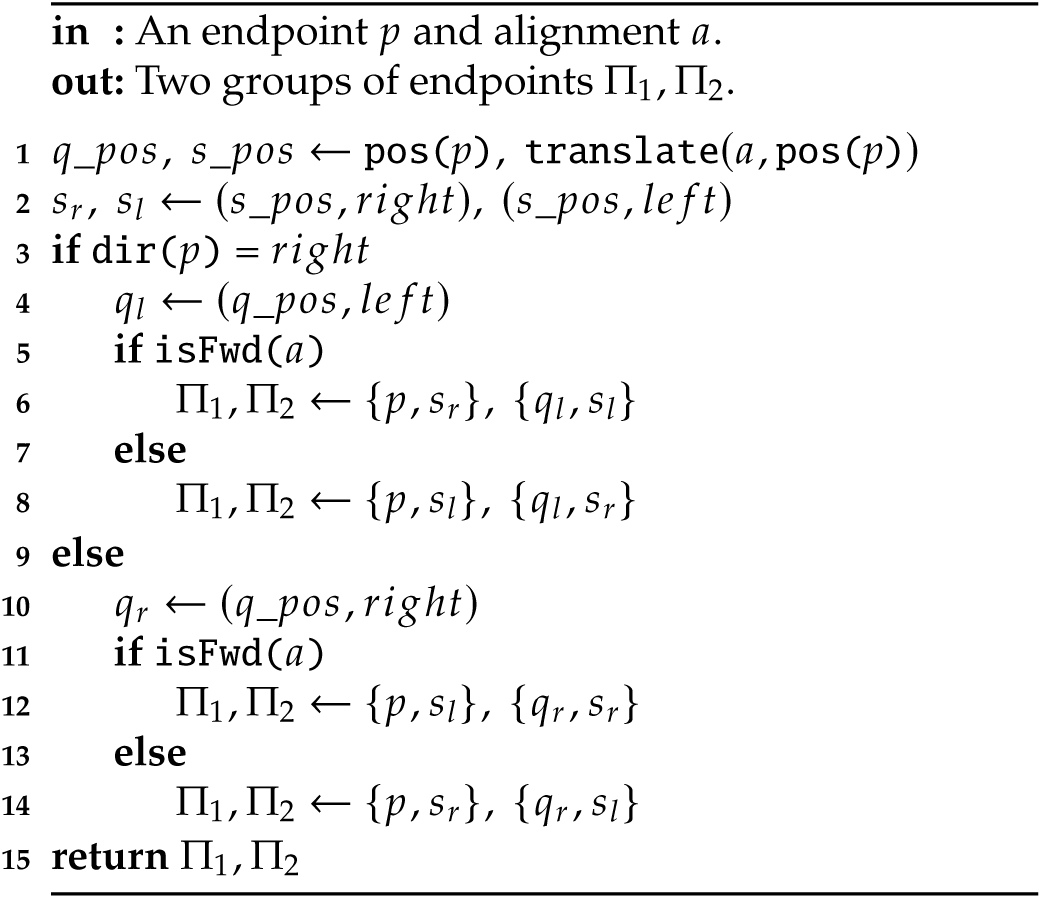

Using the consensus endpoints, we now divide each contig into disjoint intervals. We do this by constructing a set of breakpoints, ℬ, using constructBreakpoints (line 29). Each breakpoint *b* is a named tuple consisting of a position *b.*pos, and two endpoint cluster IDs, *b.*idr and *b.*idl. It occupies a single position, which marks the left and right sides of a pair of adjacent intervals that are thereby made disjoint. Iterating over each contig *c* ∈ **C** (line 2), constructBreakpoints builds two position-sorted arrays Π*_r_*and Π*_l_* which contain all consensus endpoints on *c* with *rigℎt* or *le f t* direction, respectively (lines 3-4). We then proceed through the consensus endpoints in these arrays (lines 6-7), and construct a breakpoint for each pair. If the distance between the endpoint pair is ≤ *d* (line 8), the pair is a point where two adjacent homologous intervals meet (one to the left of the pair and one to the right). In this case, we construct a breakpoint whose position is the median of the endpoints, and set *b.*idr and *b.*idl to the ID’s of the clusters that *p_r_* and *p_l_* are assigned to (line 9-12). If, however, *p_r_* is more than *d* bps to the left of *p_l_*(line 14), then the two intervals they represent do not “meet”, and the region to the left of *p_r_* is not homologous with any other region in the data. In this case, we construct a breakpoint with position set to pos(*p_r_*), *b.*idr set to clusterID(*p_r_*), and *b.*idl set to a rnd() a unique, randomly generated cluster ID (line 15). We construct a similar breakpoint in the opposite case, in which it is *p_l_* that is more than *d* bps to the left of *p_r_* (lines 17-18). Once a consensus endpoint was used to construct a breakpoint, we move to the next one by incrementing *i_r_* or *i_l_* (lines 13, 16, and 19). (Note that Π*_r_* and Π*_l_* are 0-indexed, and we define pos(Π*_x_* [*i_x_* ]) = ∞ if *i_x_* ≥ |Π*_x_* |).

### Algorithm

constructBreakpoints(𝒞, 𝒫∼)

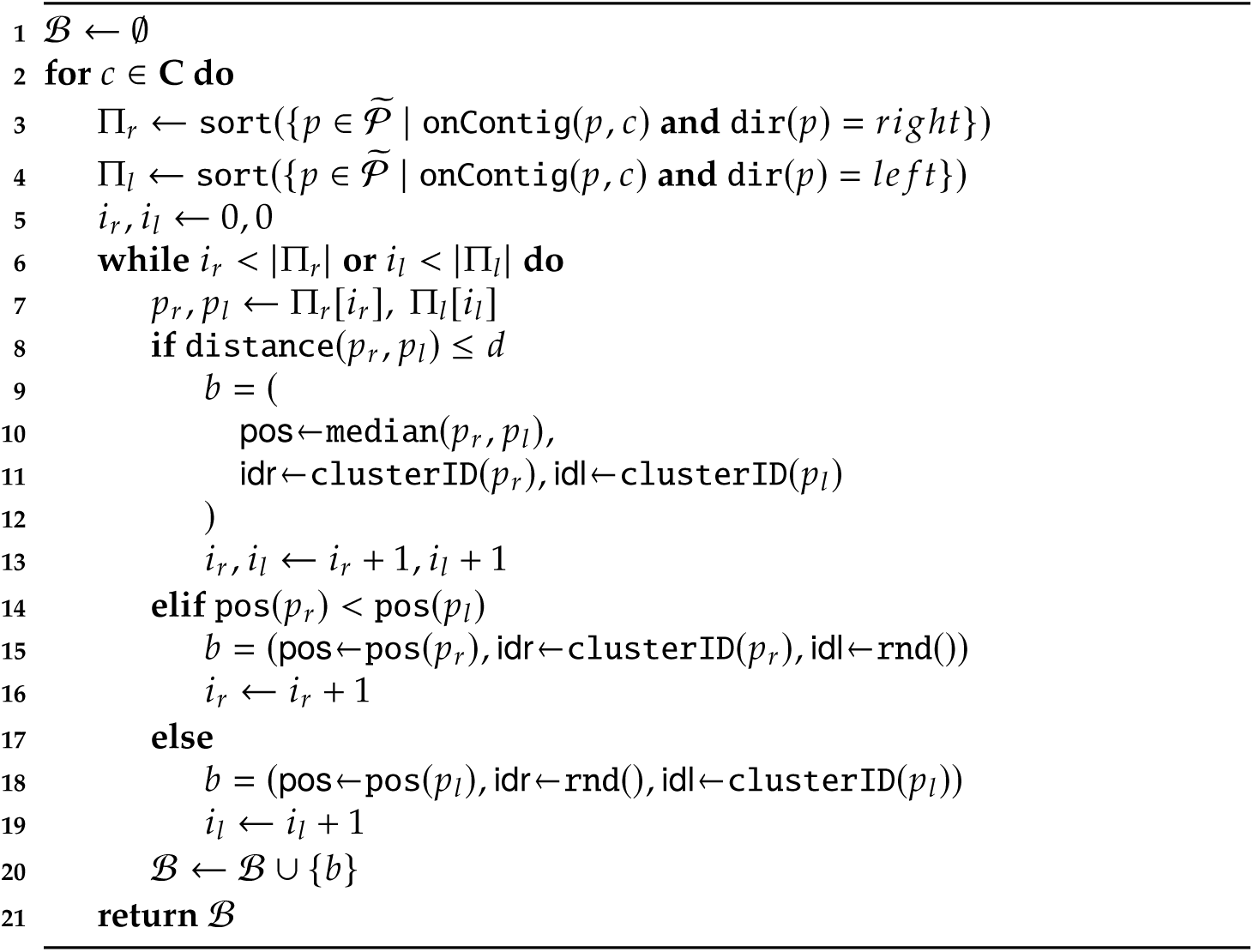

Now that we have a set of breakpoints ℬ, we can construct homology groups based on the disjoint intervals that each pair of adjacent breakpoints defines (clusterHomologyGroups, lines 30-38). Disjoint intervals, *ivl*, are defined by a named tuple that specifies both an interval on contig *c*, *c*[*ivl.*start : *ivl.*end], and a pair of cluster IDs, *ivl.*idr and *ivl.*idl, that it inherits from the breakpoints to its left and to its right, *𝐵*[*i* − 1] and *𝐵*[*i*], respectively. Note that the cluster IDs “point inwards” to the interval, i.e., *ivl.*idr is set to the *rigℎt*-referring ID of *b*[*i* − 1] and *ivl.*idl to the *le f t*-referring ID of *b*[*i*], which ensures they are set to the cluster IDs of endpoints deriving from pairwise alignments involving the interval.

Finally, we can define homology groups. Each homology group is a set of disjoint intervals, with each interval homologous to at least one other interval in the set and not homologous to any intervals outside of it. By construction, homology between intervals that make a homology group is indicated by sharing an unordered pair of cluster IDs. To show this, consider two homologous intervals, *ivl*_1_ and *ivl*_2_, which should therefore belong to the same homology group. Let *ivl*_1_ be bounded by the breakpoints *b*_1_ and 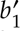, and *ivl*_2_ by *b*_2_ and 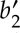, with 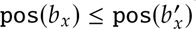. Recall that each breakpoint contains two cluster IDs, *b.*idr and *b.*idl, which refer to the regions on either side of it, and that because endpoints referring to homologous regions are clustered together, the breakpoints deriving from these endpoints share the same cluster ID. Consequently, if *ivl*_1_ and *ivl*_2_ are homologous in the same complement, *b*_1_.idr = *b*_2_.idr and 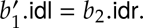, and if the intervals are homologous in the opposite complement, 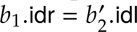 and 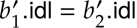. Therefore, using the equivalence between unordered pairs of cluster IDs as an indicator for homology, we assign related intervals to the same homology group. Note that following the same logic, the order within the pair of cluster IDs provides useful information, indicating which intervals are in the same complement to one another - a property we use to realize complementarity differences among intervals of a homology group in copangraph structure.

## Supplementary Algorithm 4: constructCopangraph

constructCopangraph uses the homology group information in **HGs** to merge the single-sample sequence graphs 𝒢_1:_*_𝑆_* into a multi-sample copangraph *𝑮*(**V**, **E**). The first step is to construct the copangraph node-set **V** such that each node **v** ∈ **V** represents a unique homology group. For each contig *c* ∈ **C** (line 2), we iterate over its ordered list of intervals (line 3), **HGs**[*c*]. We extract the cluster IDs *𝛼, 𝛽* of each interval (line 4) and check whether a node that represents its homology group was already created by checking its name against the unordered set {*𝛼, 𝛽*} (line 5). If not, we add a node to **V** representing the homology group (line 6). When constructing this new node **v**, we define the ordered pair of cluster IDs, (*𝛼, 𝛽*), as the ’+’ orientation by setting **v**.plus_ori to (*𝛼, 𝛽*). We then extract the node **v**, representing the homology group {*𝛼, 𝛽*} from **V** (line 7), and assign the interval represented by *ivl* to **v**, along with its sample origin and orientation (lines 8-11), which is either ’+’ or ’−’ according to the order of the cluster IDs (see **Supplementary Algorithm 3**). Consequently, each copangraph node represents all the intervals from the same homology group, along with information about their relative orientation and sample of origin.

### Algorithm

constructCopangraph(HGs, C, ℰ1:𝑆)

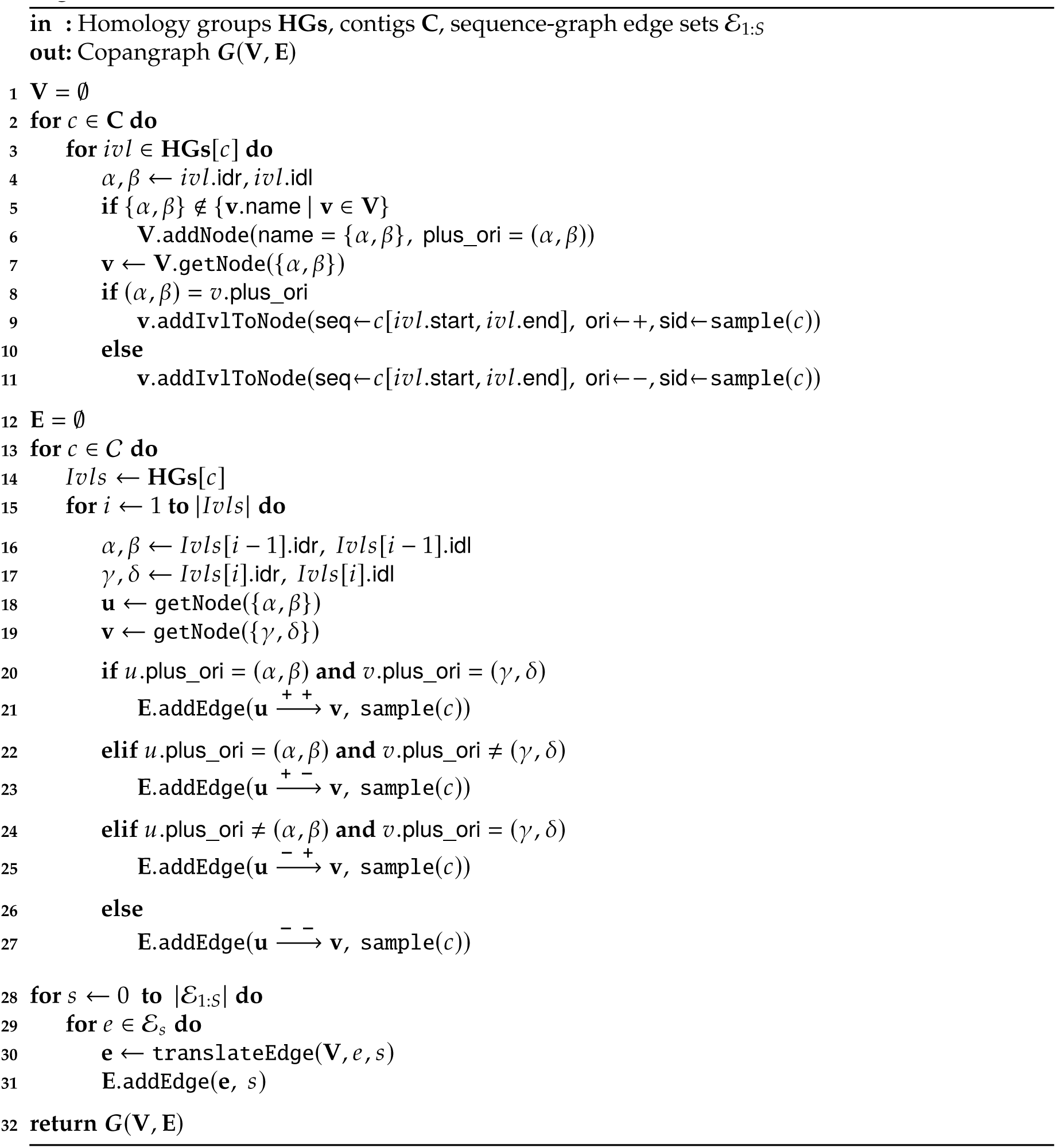

The next step is to construct the copangraph edge-set **E**. We begin by adding edges induced by contiguity between disjoint intervals (lines 12-27). For each contig *c* ∈ **C** (line 13), we extract its ordered list of intervals, and iterate over them in pairs (line 14-17). For each pair, *Ivl*[*i* − 1] and *Ivl*[*i*], we then extract the nodes **u**, **v** ∈ **V** representing the homology groups of each (lines 16-19). An edge between **u**, **v** represents the contiguity of their respective intervals in contig *c*. Edges in copangraph are bi-directed, so when adding the edge between **u** and **v** to the edge-set (lines 20-27), we set the bi-direction of the edge to the relative orientation of *Ivl*[*i* − 1] in **u** and *Ivl*[*i*] in **v**, respectively. Each edge in copangraph contains a list of samples the edge occurs in. If a particular bi-directed edge between **u** and **v** already exists in **E**, then **E**.addEdge updates the samples list of the edge, rather than duplicating the edge itself. By letting bi-direction in copangraph encode the relative orientation of intervals, complex rearrangement structure can be expressed in the graph structure and compared across samples.

Next, we add the edges of each single-sample sequence graph to the edge set **E**. We need to address two challenges: first, sequence graph edges are between contigs, but since a contig may be partitioned across multiple copangraph nodes, the correct pair of nodes should be identified and connected; and second, because single-sample sequence graphs are constructed independently from the copangraph, the bi-directed edges in each graph are not “synchronized” in their orientation. To address these challenges, we define the function translateEdge, which translates each edge from a single-sample graph into the correct copangraph edge (lines 28-31).

### Algorithm

translateEdge 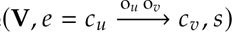

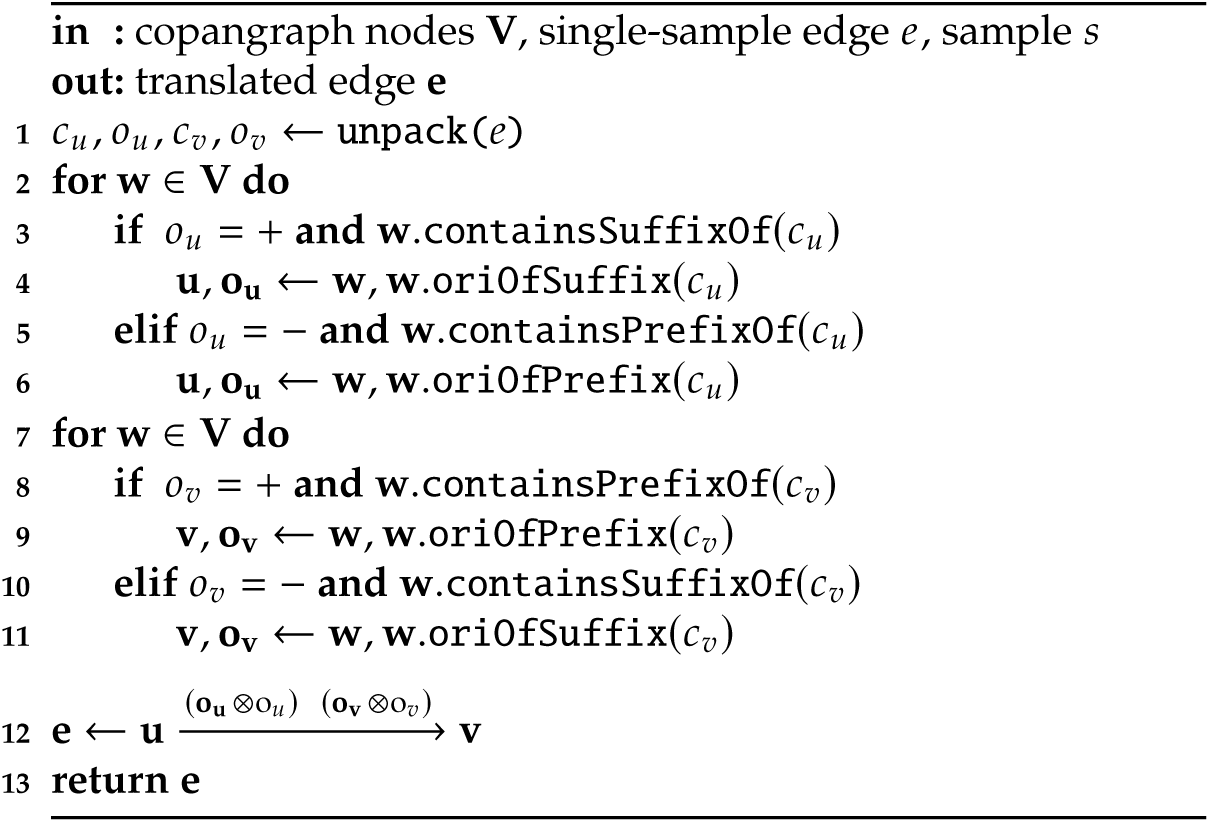

translateEdge takes an edge 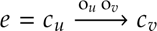 from a single-sample sequence graph and unpacks it into the pair of contigs *c_u_, c_v_* ∈ C and bi-direction information *o_u_, o_v_* ∈ {+, −} that comprise it (line 1). We then solve the first challenge, determining which copangraph nodes should be connected to translate *e* (lines 2-11). This is done by identifying the copangraph nodes containing the prefix or suffix of *c_u_* and *c_v_* based on information from the bi-directed edge *e*. Beginning with *c_u_*, which is at the tail of the arrow, we iterate through the copangraph nodes (line 2) and, if *o_u_* = +, set **u** to the copangraph node containing the suffix of *c_u_* and set **o_u_** to its orientation that node (lines 3-4). However, if *o_u_* = −, we set **u** to the node containing the prefix of *c_u_* and **o_u_** its orientation in that node (lines 5-6). We then perform a similar process for *c_v_*, which is at the head of the arrow, this time finding the copangraph node containing the prefix of *c_v_* (and its associated orientation) if *o_u_* = +, and the node (and orientation) containing the suffix of *c_v_* if *o_v_* = − (lines 7-11). We use the orientation of the prefix and suffix of *c_u_* and *c_v_* in copangraph as a reference to which we synchronize the single-sample bi-directed edges. To synchronize and translate the edge *e*, we construct a new edge **e** between **u** and **v** and combine the single-sample bi-directed information, *o_u_* and *o_v_*, with the globally consistent copangraph orientations of the intervals, **o_u_** and **o_v_**(line 12). Letting *o*_1_*, o*_2_ ∈ {+, −} we define (*o*_1_ ⊗ *o*_2_) ∈ {+, −} to be ’+’ if *o*_1_
= *o*_2_ and ’−’ otherwise. We then return the translated edge **e** (line 5) to be added to the copangraph edge-set.

